# Developmental plasticity and variability in the formation of egg-spots, a pigmentation ornament in the cichlid *Astatotilapia calliptera*

**DOI:** 10.1101/2023.04.06.535385

**Authors:** Bethan Clark, Aaron Hickey, Bettina Fischer, Joel Elkin, M. Emília Santos

**Affiliations:** Department of Zoology, University of Cambridge, United Kingdom; Department of Genetics, University of Cambridge, United Kingdom

## Abstract

Vertebrate pigmentation patterns are highly diverse, yet we have a limited understanding of how evolutionary changes to genetic, cellular, and developmental mechanisms generate variation. To address this, we examine the formation of a sexually-selected male ornament exhibiting inter- and intra-specific variation, the egg-spot pattern, consisting of circular yellow-orange markings on the male anal fins of haplochromine cichlid fishes. We focus on *Astatotilapia calliptera*, the ancestor-type species of the Malawi cichlid adaptive radiation of over 850 species. We identify a key role for iridophores in initialising egg-spot aggregations composed of iridophore-xanthophore associations. Despite adult sexual dimorphism, aggregations initially form in both males and females, with development only diverging between the sexes at later stages. Unexpectedly, we found that the timing of egg-spot initialisation is plastic. The earlier individuals establish their own territory the earlier the aggregations form, with iridophores being the cell type that responds to social conditions. Furthermore, we observe apparent competitive interactions between adjacent egg-spot aggregations, which strongly suggests that egg-spot patterning results mostly from cell-autonomous cellular interactions. Together, these results demonstrate that *A. calliptera* egg-spot development is an exciting model for investigating pigment pattern formation at the cellular level in a system with developmental plasticity, sexual dimorphism, and intra-specific variation. As *A. calliptera* represents the ancestral bauplan for egg-spots, these findings provide a baseline for informed comparisons across the incredibly diverse Malawi cichlid radiation.

## INTRODUCTION

Understanding how evolutionary changes in genetic, cellular and developmental mechanisms generate variation in organismal adult form is a fundamental goal of evolutionary developmental biology (Carroll, 2008; Smith et al., 2018). Vertebrate pigment patterns are excellent traits in which to address such questions, being extremely diverse both within and between species. Importantly, colour patterns influence a wide range of ecological interactions and are vital for animal adaptation as they can affect courtship behaviour, mate preference, predator avoidance and thermoregulation, among others (Elkin et al., 2022).

Among vertebrates, teleost fish are especially suited for elucidating the range of cellular and developmental mechanisms underlying pattern formation, since they show a strikingly diverse array of pigmentation phenotypes and the cells that generate such patterns are readily visible in the skin throughout embryonic and juvenile development (Patterson and Parichy, 2019). Yet until now, we have little insight into the cellular and developmental processes underlying teleost colour pattern formation and more specifically its variation, with most of the data coming from a few model species, such as the zebrafish (*Danio rerio*) and medaka (*Oryzia latipes*) (Irion and Nüsslein-Volhard, 2019; Kelsh et al., 2009; Owen et al., 2021; Parichy, 2021; Parichy and Liang, 2021; Patterson and Parichy, 2019).

Data from these model species show that teleost pigmentation patterns result from different proportions and arrangements of specialised colour-bearing cells, chromatophores, which synthesise and store pigments or contain reflective nanostructures. Common pigmented chromatophore classes are melanophores (black-brown pigment, melanin) and xanthophores (yellow-orange carotenoid and pteridine pigments). Rarer classes include erythrophores (red) and two classes of leucophores (white) (Parichy, 2021; Schartl et al., 2016). The structural chromatophore is the reflective iridophore, of which in zebrafish there are two distinct classes that arise by *in-situ* differentiation, appearing silver or blue depending on the structure of the refracting intracellular crystals (Gur et al., 2020).

For periodic patterns such as zebrafish stripes, the adult colour pattern is determined by appropriate behaviours of chromatophores in response to interactions between pigment cells and their surrounding tissues. These pattern formation mechanisms are best-studied in zebrafish stripe development, in which dark stripes consist of melanophores and blue iridophores while the light interstripes consist of xanthophores and silver iridophores (Patterson and Parichy, 2019). In this system, iridophores respond to early positional information provided by the tissue environment: adult iridophores first appear near the horizontal myoseptum, creating a primary inter-stripe by extensive proliferation. Subsequent stripe and inter-stripe formation is self-organising, emerging from interactions between iridophores, xanthophores, and melanophores that influence the differentiation, migration, proliferation, and death of each chromatophore (Owen et al., 2020; Volkening, 2020).

Despite the advantage of such detailed mechanistic studies, the focus for pigmentation development has been on naturally selected traits and invariant periodic patterns (such as the horizontal stripes of the zebrafish). However, many fish display colour patterns that are non-periodic, intra-specifically variable, and sexually dimorphic, such as distinctive male ornaments in for example guppies, cichlids and damselfish (Morris et al., 2020; Santos et al., 2014; Wacker et al., 2016). For such ornaments, positional information cues are hypothesised to outweigh the role of self-organising interactions, unlike periodic patterns (Parichy, 2021). Yet the mechanisms underlying non-periodic patterns and the emergence of sexually-dimorphic colouration remain unaddressed. Here we study the development of a sexually selected and sexually-dimorphic colouration ornament that exhibits inter- and intra-specific variation: the egg-spots of East African haplochromine cichlid fishes (Salzburger et al., 2007; Santos et al., 2014).

East African cichlid fish represent one of the most extensive adaptive radiations among vertebrates (Kocher, 2004; Salzburger, 2018; Santos and Salzburger, 2012; Santos et al., 2023). Their remarkable pigment pattern diversity combined with low genomic sequence divergence are an ideal system to study diversification of pigmentation traits (Malinsky et al., 2018). As a consequence, East African cichlids are increasingly used to study the genetic basis of variable colour patterns, such as stripes, bars, blotches, nuptial colouration, and also egg-spots (Albertson et al., 2014; Gerwin et al., 2021; Hendrick et al., 2019; Kratochwil et al., 2018; Liang et al., 2020; Salzburger et al., 2007; Santos et al., 2014), although the underlying cellular and developmental basis of this diversity remains unknown. To address this, we focus on egg-spots, which are yellow to red circular markings on the anal fins of males (Fig. 1A) of ∼1500 haplochromine cichlid species, varying in number, sizes, combined area, colour, and position both within and between species (Salzburger et al., 2007; Santos et al., 2014). As such, egg-spots are best described as non-periodic patterns due to varying sizes of and distances between each egg-spot within a fin.

**Fig. 1.**
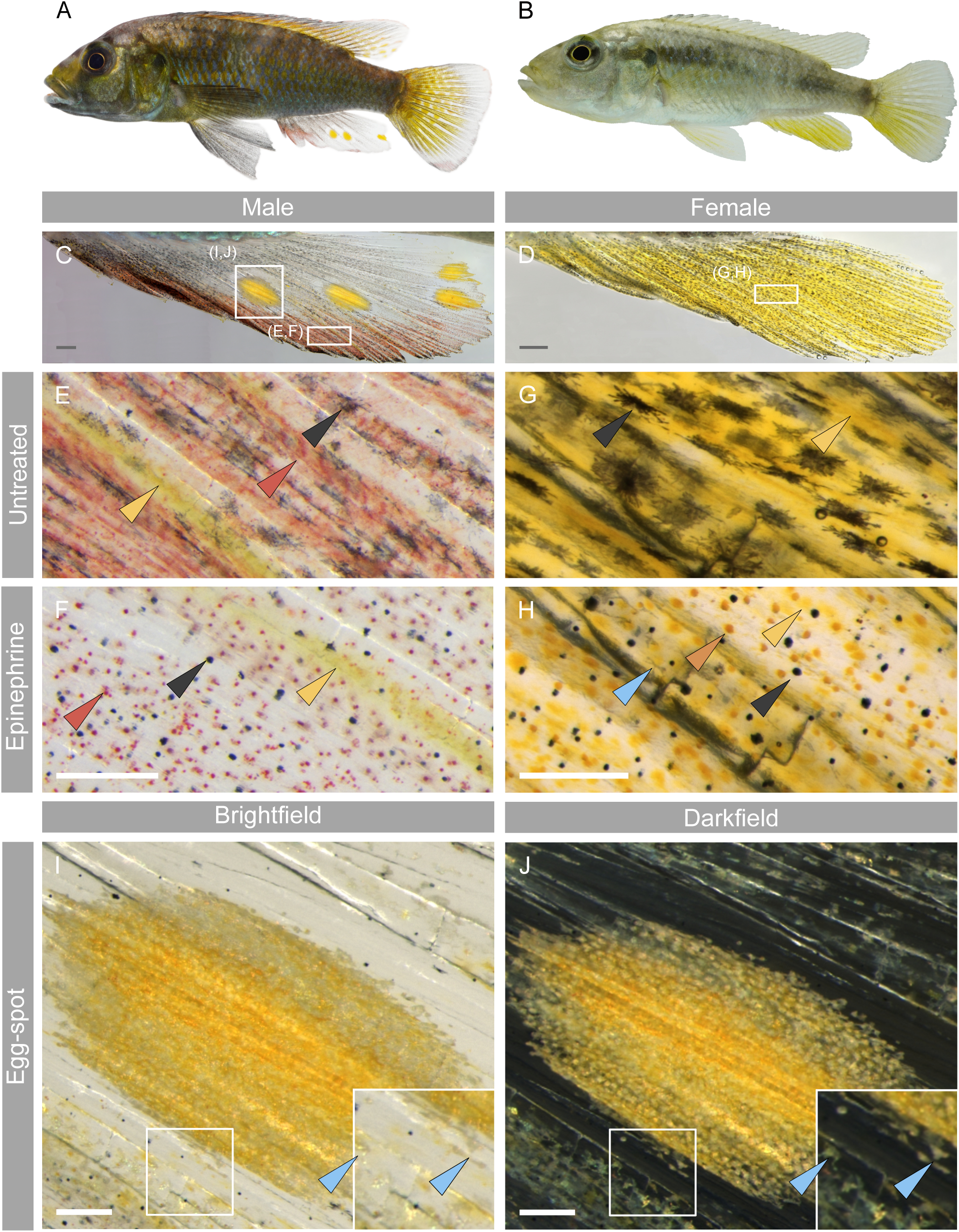
*Astatotilapia calliptera* “kisiba/masoko” male and female (deep individuals) with associated anal fin pigment cell types. Adult male (A) and adult female (B). Close up of an adult male anal fin (C) highlighting the egg-spot made up of highly packed xanthophore-like cells and iridophores. Iridophore are also sparsely distributed in other fin regions (blue arrows) (*I,J*) and other cell types present in the male anal fin such as melanophores (black arrow) and erythrophores (red arrow) (E,F). Adult female anal fin (D) highlighting cell types present throughout the fin, such as melanophores, and the two types of xanthophore-like cells, the one that contracts in response to epinephrine (orange arrow) and one other which is unresponsive to the treatment (yellow arrow) (G,H). The effect of epinephrine treatment to contract the pigment cells can be seen in (E-H). Scale bars are 1mm (C,D) and 0.25mm (E-J).

At the organismal level, egg-spots have a signalling function during the mating behaviour of these mouth-brooding fishes (Wickler, 1962). After spawning the female gathers the eggs into her mouth; the male then presents his egg-spots to which the female responds by snatching, bringing her mouth close to the genital opening that discharges sperm; the eggs then become fertilised inside the female’s mouth. The female will subsequently brood their progeny in her mouth for several weeks. Further, egg-spots also function as an honest indicator of male condition, with larger males harbouring on average a higher number of egg-spots (Lehtonen and Meyer, 2011). In some species, egg-spots are selected via female choice with females preferring males with many egg-spots over males with fewer egg-spots (Hert, 1989; Hert, 1991). In others, egg-spots have been shown to play a role in male-male competition, with males with a higher number of egg-spots having an intimidating effect on similarly sized male opponents (Theis et al., 2012; Theis et al., 2015). Taken together, these experiments demonstrated that egg-spots are sexually selected “badges of status” signalling male dominance and quality. As such, egg-spots offer a unique opportunity to study the cellular mechanisms underlying the development of intra-specific pattern variation, within the context of sexual selection.

To study the developmental basis of such pigment pattern variation, we characterise egg-spot formation in the generalist species *Astatotilapia calliptera*, which is part of the lake Malawi radiation. Phylogenetic analysis shows that all Malawi cichlid species (∼850 species) resulted from three separate cichlid radiations that stemmed from a generalist *Astatotilapia* -type ancestral lineage (Malinsky et al., 2018). As such, this species represents the ancestral bauplan of egg-spots for the Malawi cichlid radiation (Turner et al., 2021). More specifically, we focus on a population of *A. calliptera* from an isolated crater lake north of Lake Malawi referred to in the literature as Lake Masoko (variant spelling Massoko, as used by the German colonial administration) and known locally as Lake Kisiba (Clark et al., 2021; Turner et al., 2019).

The *A. calliptera* from Lake Kisiba are at an early stage of adaptive divergence into two ecomorphs – benthic (deep water) and littoral (shallow waters) (Malinsky et al., 2015). Both the “deep” and “shallow” individuals display intra-specific variation in egg-spot number and colour. Using these populations, we aimed to: 1) identify key cellular events contributing to egg-spot development and variation; 2) uncover when and how anal fin pigmentation development diverges between males and females; and 3) determine if external conditions, such as timing at which individuals became dominant, influences egg-spot development.

We find that both iridophores and xanthophore-like cells are involved in egg-spot development, with a key role for iridophore aggregations in initialising egg-spot formation. We show that both males and females exhibit initial stages of egg-spot formation, with sexual dimorphism becoming apparent only at later juvenile stages. Furthermore, we find that the timing of the onset of egg-spot development is plastic and does not correlate with age or size. Instead iridophore aggregations form sooner when the fish is isolated from conspecifics and becomes the dominant individual in its own territory. Finally, we characterise consistency in the position of the first egg-spot, variability in subsequent egg-spot positions, and features of development which may contribute to adult variation in this pattern.

We propose that cellular interactions explain much of the process of egg-spot development, suggesting that non-periodic pattern formation can be governed by similar mechanisms as periodic patterns, though we also identify cellular interactions and behaviours not observed in other systems like zebrafish stripes. In addition, we suggest multiple inputs involving positional information and hormonal mediation to regulate egg-spot position consistency, onset timing plasticity, and sex-limited maturation. Our results indicate that a combination of cellular interactions and these additional inputs could be developmental sources of variation in egg-spot patterns.

Taken together, the haplochromine cichlid egg-spots provide an exciting model system to investigate how environmental and behavioural cues affect pigment pattern formation at the cellular and developmental level. Further, *A. calliptera* represents the ancestral bauplan for egg-spots, as such our findings provide a basis for comparison of patterning mechanisms across the Malawi radiation which harbours an incredible extent of inter- and intra-specific variation in egg-spot patterns.

## RESULTS

### Male and female anal fins have different pigment cell composition and distribution

We first sought to characterise the cellular basis of sexual dimorphism in the anal fin by imaging adult fins, finding differences in the presence and arrangement of pigment cell types between males and females (Fig. 1A-H). The conspicuous egg-spots in sexually mature *A. calliptera “kisiba/masoko”* males are composed of densely packed iridescent iridophores and yellow-to-orange xanthophore-like cells (Fig. 1I-J), similar to *Astatotilapia burtoni* (Santos et al., 2014). The egg-spots’ conspicuous nature is enhanced by the presence of an outer transparent ring surrounding each egg-spot. This region appears to have a reduced amount or is devoid of pigment cells altogether (Fig. 1I-J). This transparent ring provides contrast and is specific to males (Fig. 1C,D). Xanthophore-like cell distribution is mostly concentrated within the egg-spots in adult males, while iridophores are packed within the egg-spots but are also sparsely distributed across the anal fin (Fig. 1C,I,J). Iridophore identity was confirmed due to the cell’s iridescence, which can be seen using both brightfield and dark field reflected light conditions (Fig. 1I,J), Melanophores are distributed across the anal fin, but absent in egg-spots (Fig. 1C,E,F,J). Additionally, males have red erythrophores on the distal edge of the anal fin (Fig. 1C,E,F).

In contrast, females display a homogenous distribution of xanthophore-like cells across the anal fin resulting in an overall yellow fin (Fig. 1D,G,H). Upon close inspection, we found two morphologies of xanthophore-like cells in the female fins. One type contracted in response to the epinephrine treatment, revealing a dark orange centre surrounded by a yellow halo (Fig. 1H orange arrow, Fig. S1). The other type was lighter in colour and unresponsive to epinephrine (Fig. 1H, yellow arrow, Fig. S1). This unresponsiveness persisted when using higher epinephrine concentrations and longer exposure times (Fig. S1), but whether this difference in colour and contractility represents a difference in cell type remains to be determined. The presence of two xanthophore-like morphologies in females may also suggest that such cells are also present in male egg-spots, and that perhaps we were unable to identify them due to their high density which makes imaging single cells difficult at such stages. Additionally, adult female anal fins also have melanophores and iridophores sparsely distributed across the anal fin but lack any erythrophores (Fig. 1D,G,H).

To investigate egg-spot development and the ontogenetic origin of sex specific differences, we characterised the sequence of cellular events from the onset of overt embryonic fin development to adult stages.

### Iridophore aggregations initiate egg-spot formation in both sexes

To identify the chromatophores and cell behaviours that initialise egg-spot development, we characterised the earliest visible colouration events. For this purpose, we imaged the same individuals through time from the onset of overt anal fin development to juvenile stages (Fig. 2). More specifically, we imaged: 1) the fins of embryos, single housed at 10 days post fertilisation (dpf), up to yolk closure (embryo cohort, Fig. 2A); 2) the fins of early juveniles, isolated at the stage of yolk closure, during a period of 50 days (yolk closure cohort, Fig. 2B). Two cohorts were required, as it was not possible to continue imaging embryo cohorts after yolk closure, because repeated anaesthesia as juveniles after daily anaesthesia as embryos caused low survival rates (see methods for more detail). Despite this, we found a stereotyped series of events during early anal fin pigmentation development that was consistent between individuals, replicates of different morphs, and cohorts.

**Fig. 2.**
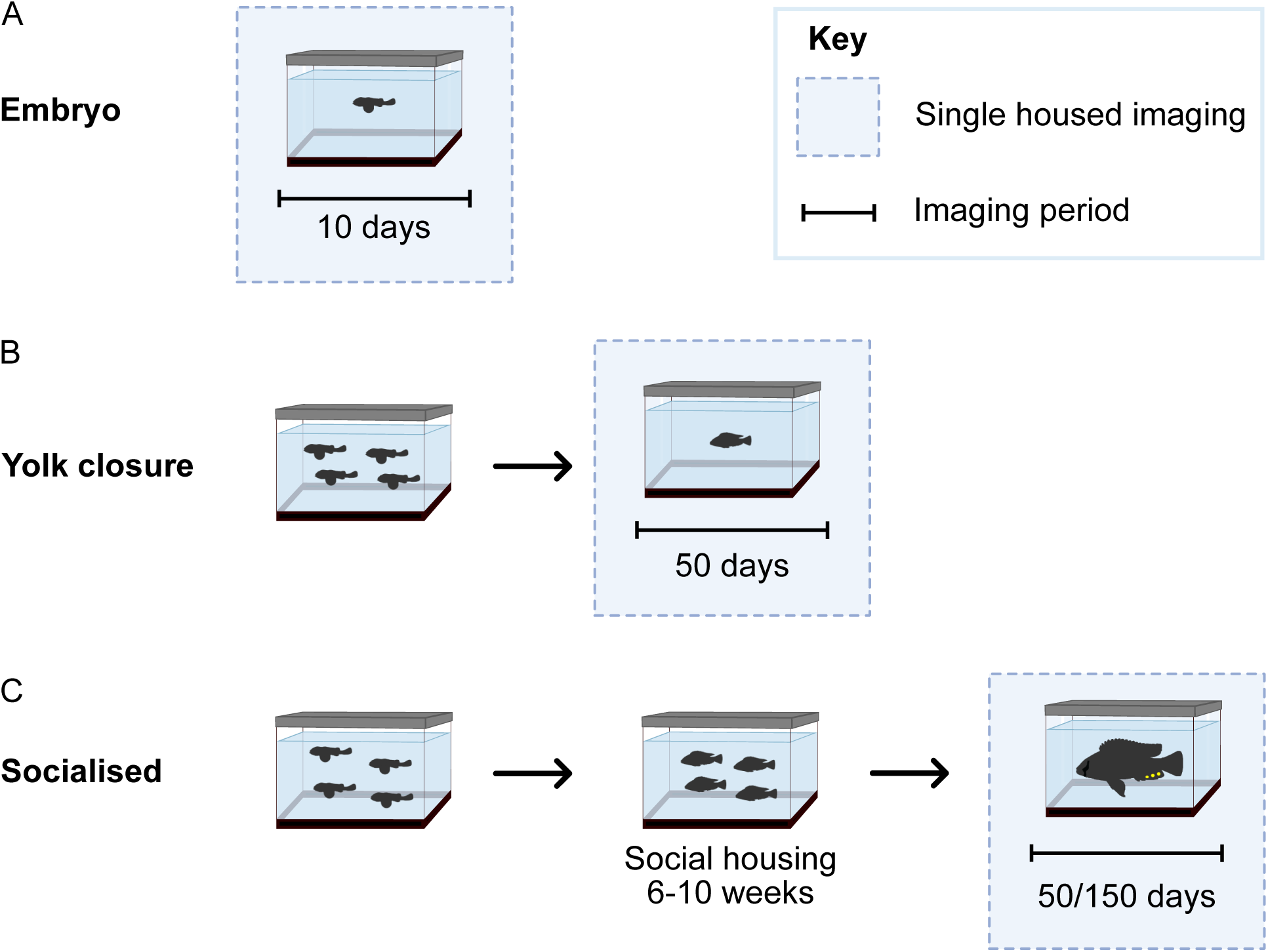
Experimental design of longitudinal imaging for the three cohorts: embryo, yolk closure and social. To characterise egg-spot development at the cellular level, we followed the same individuals through time, from the onset of overt embryonic anal fin development (10 dpf) to adult stages showing three mature egg-spots (∼220 dpf). To span all the developmental events that lead to the formation of egg-spots we performed three imaging series starting at three different developmental stages: 10 dpf embryos (A), juveniles at the point of yolk closure (B) and juveniles maintained in a social group for 6 to 10 weeks post yolk closure for deep and shallow individuals respectively (C). For each cohort stage, two replicates were completed, one with shallow individuals and one with deep individuals. All individuals were kept in a group before being housed in individual compartments/territories for imaging (B-C). Imaging period refers to the time during which each cohort was maintained in single housing; within this time period individual anal fins were repeatedly imaged every few days (see methods for details). Embryo and yolk closure cohorts span the initial stages of egg-spot development (initial aggregations of pigment cells (section 3*b*)), whereas the socialised cohorts span the latest events (transparent ring formation (section 3*c*)).

First, background pigmentation develops in the fin, with melanophores appearing in embryos followed by xanthophore-like cells (Fig. 3A,B). Melanophore and xanthophore-like cell coverage increases throughout embryonic stages (Fig. 3C,D,M). At the transition between embryonic and juvenile development (yolk closure), the anal fin has melanophores homogeneously distributed throughout the fin (Fig. 3C-F) and xanthophore-like cells spread across the anterior region of the fin up to the sixth fin ray, all contracting in response to epinephrine (Fig. 3C-F). Melanophore numbers in the fin decline in early juveniles soon after yolk closure(Fig. 3H,N,M), while contracting xanthophore-like cell coverage continues to increase (Fig. 3E-J).

**Fig. 3.**
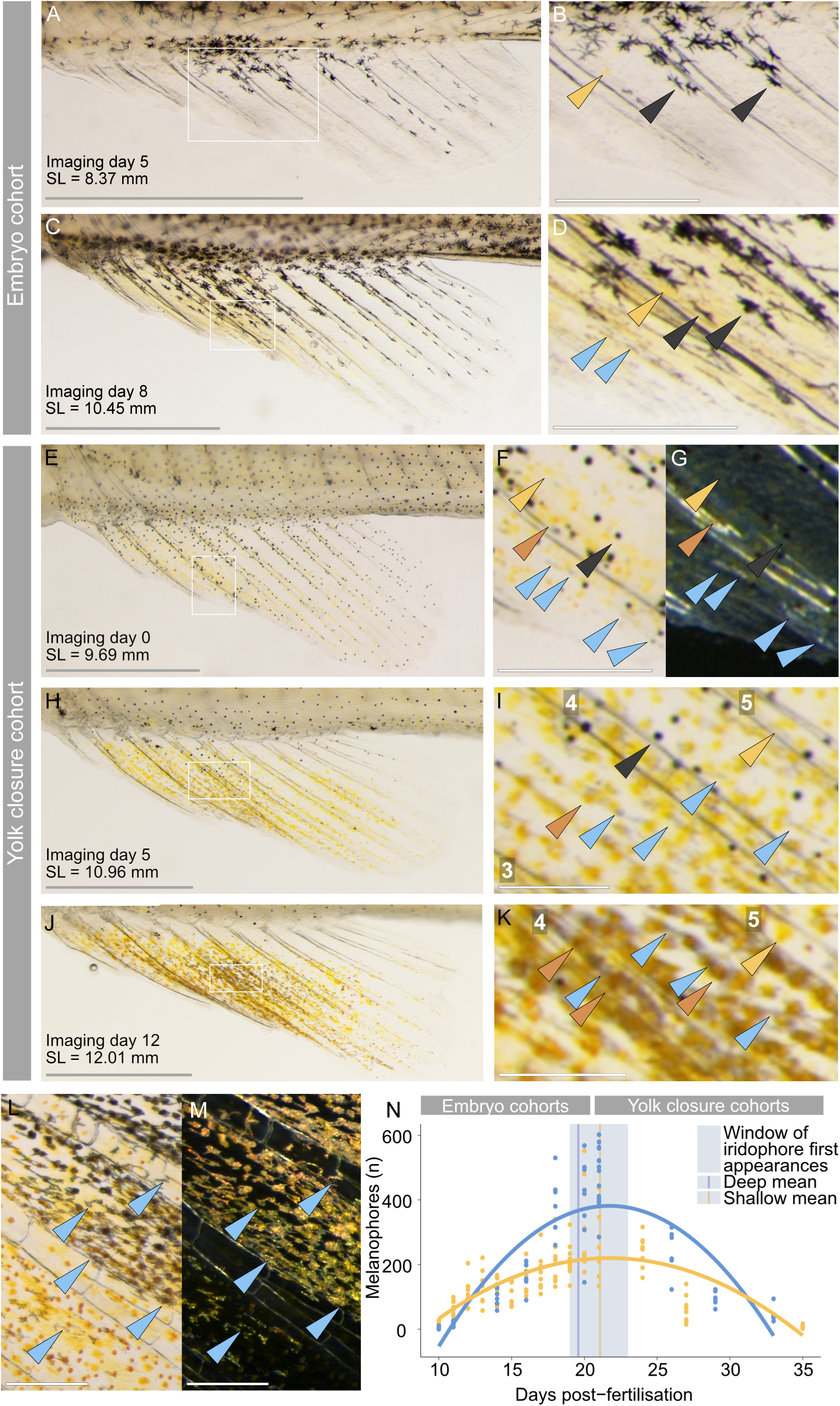
Description of early anal fin pigmentation ontogeny leading up to the initial stages of egg-spot development in *A. calliptera* “masoko/kisiba” (A-J) Images of developing fins showing the early development of anal fin pigmentation and egg-spot initialisation in embryo and yolk closure deep cohorts. All images are with transmitted light except (G) and (M) under reflected light. Events shown are: melanophores and xanthophore-like cells present in the embryo fin (A,B), spread of chromatophores in the embryo fin and appearance of iridophores (C,D) xanthophore-like cells and iridophores first present in early juvenile fin under different lighting conditions(E,F,G), iridophores first present in position of first spot, i.e. between fourth and fifth fin ray (H,I) iridophores associate with xanthophore-like cells (J,K). (L-M) Iridophores associated with contracting xanthophore-like cells under different lighting conditions (blue arrows). (N) Number of melanophores in the fin in embryo and early juvenile cohorts days post fertilisation, with grey shading indicating the timing in which iridophores first appeared in the fin (19-23 dpf for both shallow and deep individuals across embryo and yolk closure cohorts). Arrows indicate: melanophores (black), contracting xanthophore-like cells (orange), uncontracting xanthophore-like cells (yellow), iridophores (blue). Grey scale bars 1 mm, white scale bars 250 µm. Embryo fins are not treated with epinephrine, early juvenile fins are treated with epinephrine causing the pigments in the chromatophores to contract.

The appearance of iridophores is the first visible sign of egg-spot development. In both embryo and yolk closure cohorts, iridophores first appeared in the anterior hard-spined domain of the anal fin in the days surrounding yolk closure (19-23 dpf) (Fig. 3D,F,G,N). Iridophore appearance coincides with peak melanophore numbers in the fin (Fig. 3N). iridophores have a thin elongate appearance which appear as faint grey lines under transmitted light and reflective under reflected light (Fig. 3F,G,L,M). Iridophores first appear near the third fin ray (Fig. 3C-G) and then appear between the fourth and fifth fin rays, accumulating from 3 to 6 days after first iridophore appearance (Fig. 3H-I). Notably, the first egg-spot to develop always formed centred between the fourth and fifth fin rays (Fig. 3J-K) making the arrival of iridophores to this landmark the earliest visible event of egg-spot formation. When iridophores are present between the 4th and 5th fin rays, both uncontracting and contracting xanthophore-like cells are seen in the rest of the fin.

Next, an increasing number of iridophores accumulate between the fourth and fifth fin rays (*Fig. 3*J-K), often spreading posteriorly up to the sixth fin ray. Iridophores become so numerous and connected that individual cells are hard to distinguish (Fig. 3K-L). As their number increases, iridophores become associated with xanthophore-like cells: in epinephrine-treated fins, contracting xanthophore-like cells are visibly associated with iridophore (Fig. 3K-M). The xanthophore-like cells in these associations have a dark orange colour, and a lighter halo is either much smaller or cannot be seen, which may be due to stronger contraction, obstruction by associated iridophores, or indicate a different cell type. This association between iridophores and xanthophore-like cells is observed with high frequency and consistency but it is unclear whether the xanthophore-like cell is simply overlaying the iridophore, there is a stronger connection between the two cells in the association, or the apparent association is a distinct differentiated form of iridophores containing pigment as well as refractive crystals.

The described order of developmental events that contribute to the initiation of egg-spot development is consistent between individuals and replicates of different morphs, with greater timing variation for later events, whether measured by days since initial imaging session, standard length (SL, a size indicator of developmental stage and nutrition-related condition), or SL growth since initial imaging session (Fig. S2 and S3). Taken together, these results suggest that iridophores have a role in initialising egg-spot formation even though they are not the first chromatophores to appear in the fin. The initialisation by iridophores is followed by the involvement of xanthophore-like cells, while melanophores and erythrophores do not seem to play a role in these initial stages, which is consistent with the cellular composition of adult egg-spots. Surprisingly, both males and females show the same developmental trajectory in these early stages, as all individuals develop iridophore aggregations, with no difference in order of events or growth rate (Fig. S4).

### Transparent ring formation is concomitant with a shift in xanthophore density which occurs only in males

At the end of the imaging of the yolk closure cohorts (50 days post yolk closure) (Fig. 2B), all individuals displayed large iridophore and xanthophore aggregations, but lacked distinctive features of the adult male egg-spot pattern, such as the chromatophore-free transparent ring surrounding each egg-spot (Fig. 1A,C,I,J). Furthermore, there were no visible differences between male and female anal fins at this stage, despite mature egg-spots being a sexually-dimorphic trait in *A. calliptera*. To determine the cellular events leading to the formation of mature egg-spots and to characterise pigmentation development divergence between the sexes, we imaged anal fin pigmentation development in late juveniles. These individuals were initially kept together in a typical social stock tank (socialised cohort), and were transferred to single housing for imaging ∼6 to 10 weeks after yolk closure. We followed two socialised cohorts: shallow individuals isolated after ∼10 weeks showing early aggregations followed for 50 days, and deep individuals isolated after ∼6 weeks followed for 150 days (Fig. 2; see methods for details).

The socialised cohort exhibited the same consistent order of events leading up to egg-spot initialisation (Figs. S5 and S6). Similarly to yolk closure cohorts, the initial aggregations formed in socialised cohorts contain iridophore-xanthophore associations and are similar between males and females (Figs. 4A,F and S7). Subsequently in all males in the socialised cohort, xanthophore coverage decreased outside the egg-spot region (Figs. 4A-C, I *and S7*) while female xanthophore distribution across the fin remained stable throughout the study (Figs. 5F-I *and S7*). We found a significant inverse correlation between xanthophore-like cell area coverage outside of the egg-spot region and day since first imaging session in males (Pearson’s correlation R = −0.75, P<0.0001), while no correlation was found in females (Pearson’s correlation R = −0.09, P = 0.53). Concurrent with loss of xanthophore coverage, the iridophore-xanthophore aggregations become increasingly dense in males (Figs. 4A-C and S7) but not in females (Figs. 4F-H and S7). As aggregations grow, very dark contracting xanthophore-like cells similar to those in egg-spots appear elsewhere in the fin, without association with iridophores, in addition to the lighter contracting and uncontracting xanthophores, but only persist in females (data not shown). The appearance of the transparent ring was the final major event in egg-spot formation (Fig. 4C) and is similarly limited to males, coinciding with reaching a near-total loss of xanthophore-like cells outside of the egg-spot region (Figs. 4C,I and S7). As the transparent ring forms, melanophores and iridophores appear outside the egg-spots, sparsely and homogeneously distributed throughout the fin without associating with other pigment cell types present (Fig. 4D-E). Additionally red erythrophores appear at the distal edges of the fin only in male *A. calliptera* (Figs. 4D-Eand S7).

**Fig. 4.**
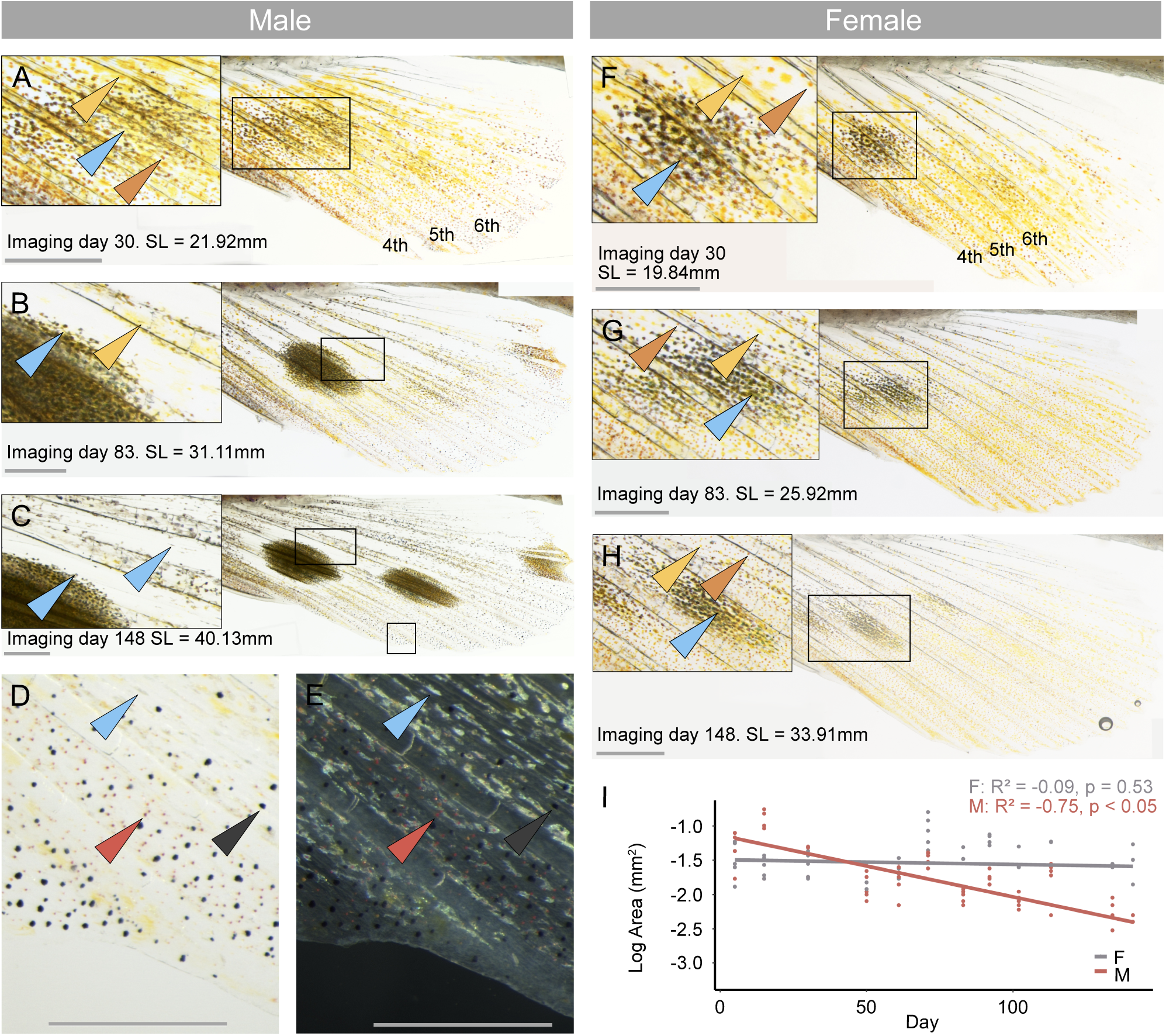
Divergence in egg-spot development between sexes of *A. calliptera* “masoko/kisiba” deep individuals. Images indicate shared development (A,F) and point of divergence (B,G) in juvenile males and females. Insets highlight the first egg-spot aggregation region between finrays 4-6. Points of divergence can be seen from the reduction of xanthophore distribution across the fin in males (B) unchanging in females (G) and the appearance of a transparent ring surrounding mature egg-spots (C) absent in females (H). Iridophores, erythrophores and melanophores (D-E) are found in mature males (C) outside of the egg-spot region. (I) Area coverage of xanthophore-like cells (both contracting and uncontracting) was measured in a 0.5mm^2^ square region centred on the first segment of the eighth fin ray. Male xanthophore-like area coverage reduces throughout the study whereas it remains constant in females. Arrows indicate: melanophores (black), contracting xanthophore-like cells (orange), uncontracting xanthophore-like cells (yellow), erythrophores (red) and iridophores (blue).

**Fig. 5.**
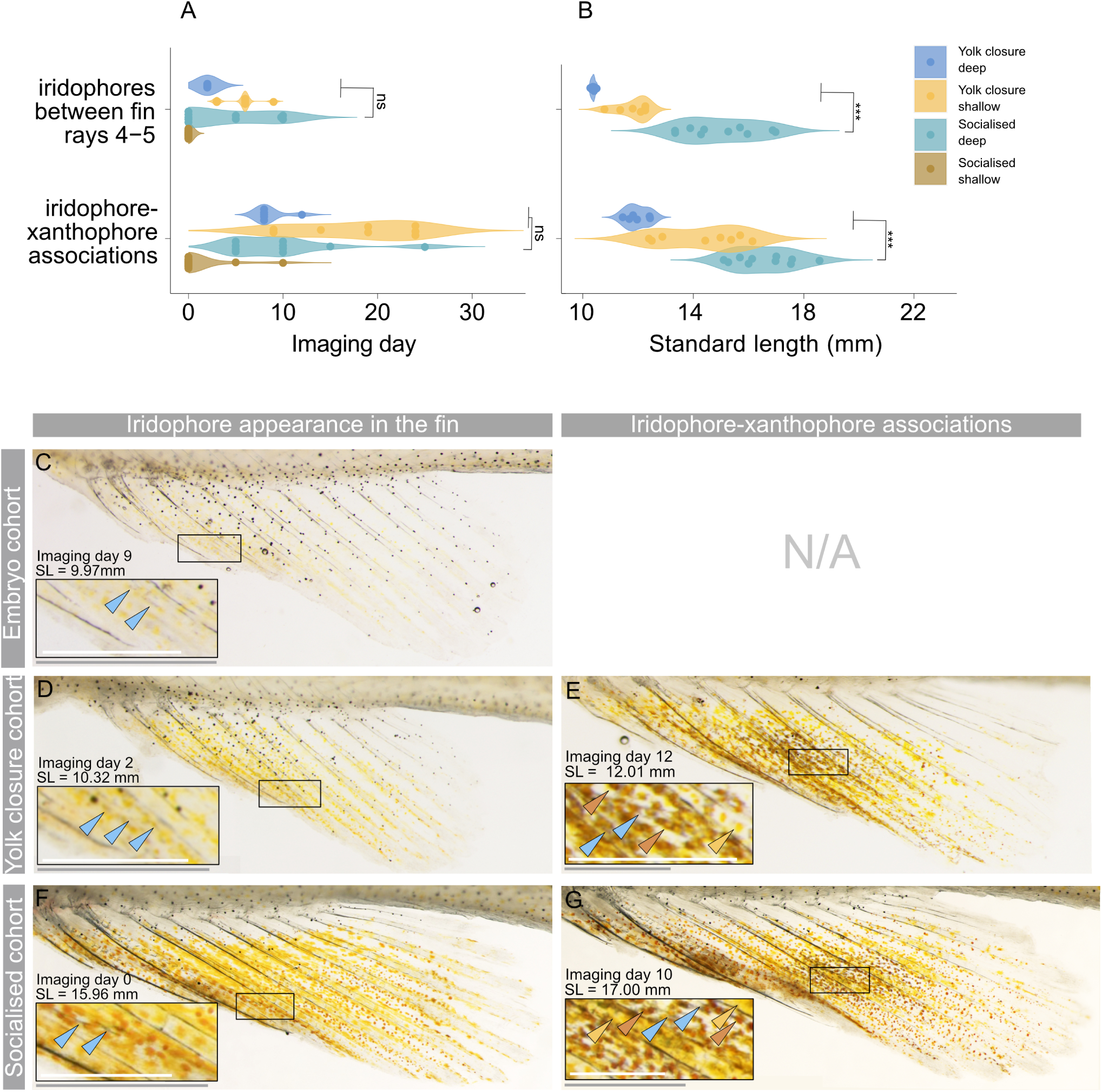
Iridophores aggregate soon after fish are singly housed at different developmental stages, despite differences in size of fish and development of background pigmentation. (A-B) Violin plots of iridophore events in early egg-spot formation by cohort and morph. For the first imaging session in which the event was observed, each event is plotted against days since the first imaging session (Day) (A), and standard length (SL) (B). Each point is one individual. Standard length measurements were not obtained for the socialised shallow cohort. (C,D,F) Transmitted light images of the fin when iridophores are first present, for a representative deep morph individual from the embryo cohort (C), yolk closure cohort (D), and socialised cohort (F). (E,G) Transmitted light images of the fin when iridophore-xanthophore associations are formed, for a representative deep morph individual from the yolk closure cohort (E), and socialised cohort (G). Associations do not form before the end of embryo stages. Arrows indicate melanophores (black), xanthophores (orange) and iridophores (blue). Grey scale bars 1 mm, white scale bars 250 µm.

These results show a common developmental program of cellular behaviour between the sexes from embryonic stages to early juvenile stages, where associations between iridophores and xanthophores form, followed by divergence in late juvenile stages. More specifically, in females aggregation growth plateaus and xanthophore coverage elsewhere in the fin persists. Females at the end of our series continue to display small aggregations despite their absence in most female adults (Fig. 1B). Further imaging of later stages would be needed to characterise how and when these aggregations are lost. Only in males do aggregations mature into egg-spots, by continuing to increase in size and density with decreasing xanthophore coverage elsewhere in the fin, followed by the formation of the transparent ring. Therefore the earliest visible point of divergence between sexes is the concordant growth of aggregations and reduction in xanthophore-like cell distribution across the fin in males (Fig. 4B,G).

### The onset of egg-spot development is plastic and dependent on social conditions

The timing at which an individual becomes territorial and hence dominant may impact pigmentation development through hormonal control (Maan and Sefc, 2013). Dominant cichlids rapidly become more colourful, thus we sought to test if environmental factors such as timing at which *A. calliptera* individuals are single housed and gain their own territory influences any aspect of egg-spot development. To test this, we took advantage of the fact that our three imaging series started at three different developmental timepoints (10 dpf embryo, yolk closure juvenile and late juvenile; see Fig. 2), with individuals being singly housed at different ages. For each event in egg-spot initialisation (xanthophore appearance, iridophore appearance, iridophores position in 4^th^ & 5^th^ fin ray, and iridophore-xanthophore associations), we contrasted the number of days since the first imaging session and standard length. The number of days reflects the elapsed time since the individual was single housed, and standard length is a descriptor of developmental stage which is a preferred indicator over dpf (Parichy et al., 2009; Singleman and Holtzman, 2014) (Fig. S8). We reasoned that if there is plasticity in egg-spot development induced by the timing at which individuals are single housed, then for such events, days since single housing would be a better predictor of event timing than standard length (developmental stage).

The timing of the two first events, appearance of xanthophores and iridophores in the fin, is better predicted by developmental stage than time in single housing (Fig. S9). The first appearance of xanthophores is always during embryo stages (Fig. S9, Table S2) and the first appearance of iridophores in the fin is around yolk closure (Fig. S9, Table S2) - between one day prior and three days after yolk closure (Fig. 3N and Fig. 5C-D) causing iridophores to be visible in the fin typically from the first imaging day in the socialised cohort (Fig. 5F, S6, S9). However, after the first appearance of each pigment cell, only xanthophore-based pigmentation progressed with developmental age. Consequently, on the first day iridophores are observed in the fin for each cohort, the appearance of iridophores is similar between cohorts, with very few cells in the anterior of the fin (Fig. 5C,D,F). Meanwhile, the xanthophore-like coverage is lowest in the embryo cohort (Fig. 5C), greater in the yolk closure cohort (Fig. 5D) and greatest in the socialised cohort with both contracting and uncontracting morphologies visible (Fig. 5E).

In contrast, changes in iridophore-based pigmentation in the fin first occur only in the days after single-housing regardless of each cohort’s developmental stage. Iridophores appeared between the 4th and 5th fin rays 0-9 days after single-housing in yolk closure cohorts (20-29 dpf) and 0-10 days after single-housing in the deep socialised cohort (∼42-52 dpf) (Fig. 5A,E,G). Associations appeared 8-24 days after single-housing in yolk closure cohorts (28-44 dpf) and 5-25 days after single-housing in the deep socialised cohort (∼47-67 dpf) (Fig. 5A,E,G). Only in the shallow socialised cohort were iridophores present between the 4th and 5th fin rays and associations present on the first day of imaging (n=12 out of 12, n=10 out of 12 respectively) (Fig. 5A). This was due to the shallow socialised cohort being singly-housed at the latest stage of all cohorts, with individuals selected for visibly initiated egg-spot development (existing aggregations) (see methods). Together, this demonstrates that when raised in a group, egg-spot development involving iridophores only progressed beyond appearance in the fin at later stages, but can occur sooner upon earlier gain of territory.

Consequently, when socialised shallow individuals are excluded due the bias towards existing aggregations, the standard lengths at the point of these later iridophore events are significantly different between yolk closure and socialised cohorts(Fig. 5B, Table S2), but the days since initial imaging sessions are not significantly different (Fig. 5A, Table S2). Days in single housing are therefore a better predictor of iridophore-based egg-spot formation events than developmental stages. Furthermore, the duration of egg-spot initialisation was unaffected by the developmental stage at which isolation commenced (Table S2).

We infer that individual isolation can induce aggregation development, regardless of developmental stage and progress of fin background pigmentation. This indicates that the initiation of iridophore aggregation is condition-dependent and that the progression of xanthophore-based background fin colouration is independent from social condition, until it responds to iridophore events (see Fig. 4). As aggregation development is initiated by the appearance of iridophores in the position of the future spot (Fig. 3), we suggest that iridophores are the social condition-dependent cell type which cause a change in the fin colouration development thitherto.

### Competitive interactions between aggregations likely contribute to egg-spot pattern variation

In *A. calliptera,* there is variation in the number of egg-spots on the anal fin. Egg-spots continue to be added as juveniles and adults continue to grow, so that typical adult spot numbers in *A. calliptera “masoko/kisiba”* range from 2 to 17 (*unpublished data*). Though the initiation timing of the first egg-spot is condition-dependent, the process of egg-spot formation is consistent between individuals for the first egg-spot. We asked whether other aspects of egg-spot development could facilitate egg-spot number variation, and how early this variation arises. For this purpose, we examined variation in the development of the first three egg-spots in males in our socialised cohorts.

Unlike the first egg-spot, the formation of subsequent egg-spots is dynamic and variable between individuals. Multiple aggregations may initially form adjacent to each other, separated by single fin rays (Fig. 6A, white arrows), or more distant to other aggregations, separated by multiple fin rays (Fig. 6B, white arrows). In some fins, the first aggregation is initially large and spans multiple fin rays but in these cases, this first aggregation reduces in spread while increasing in density, so that it does not cover the posterior regions of the fin where subsequent aggregations later appear (Fig. 6*C*, white arrows).

**Fig. 6.**
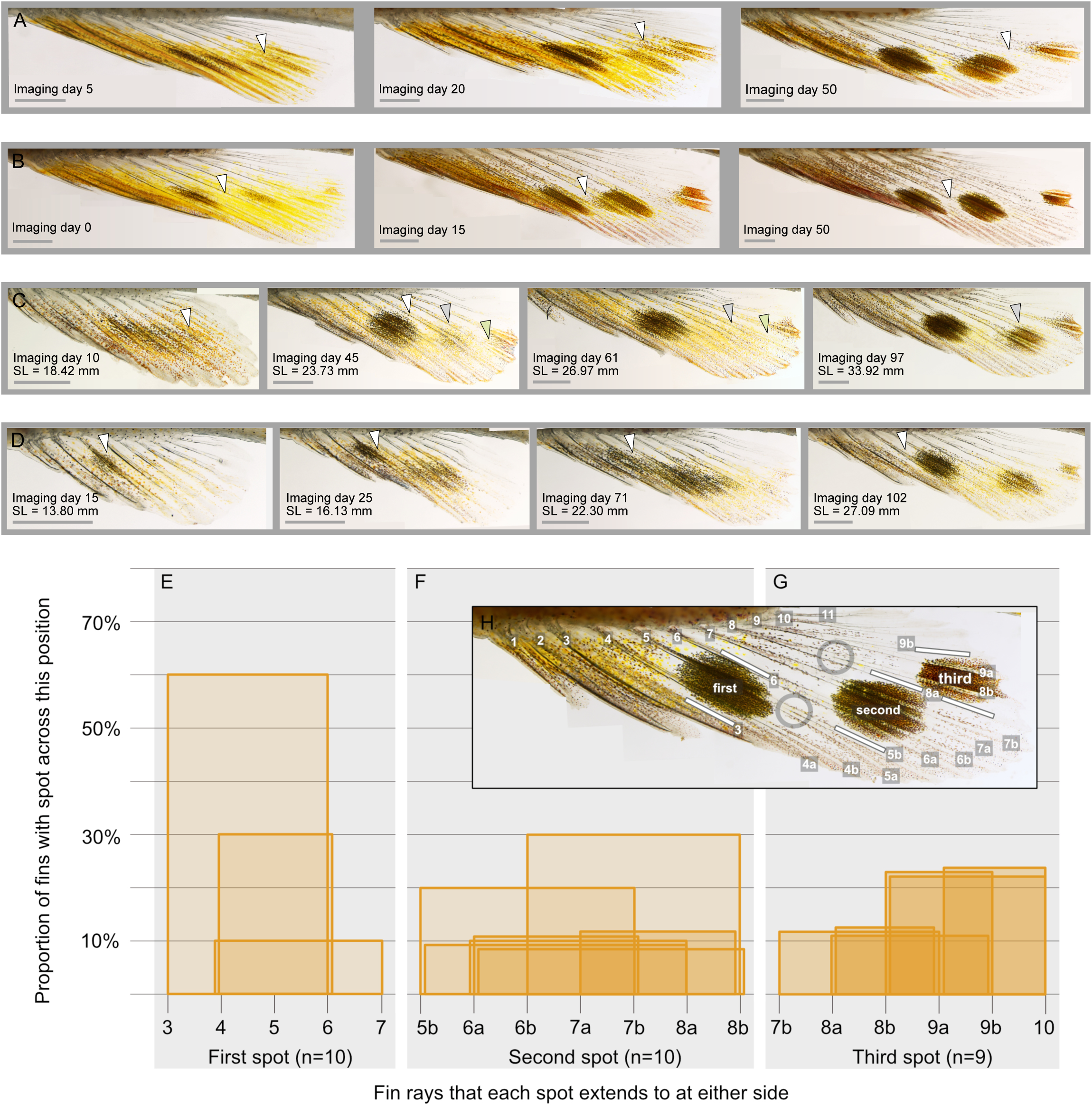
Variability and flexibility in subsequent egg-spot formation. (A-D) Examples of dynamic formation of second and third egg-spots. Arrows indicate points of interest for emerging and fading aggregations. Each row is one male from a socialised cohort. Grey scale bars 1 mm. (E-G) Proportion of male individuals from socialised cohorts with an egg-spot at each position for the 1st (E), 2nd (F), and 3rd (G) egg-spots, of those that have each egg-spot. Positions are described by the fin rays that the aggregations are bounded by at either side, measured on the final imaging session, counting only mature egg-spots with dense and defined cellular aggregations. Each of the three initial spots are diagrammed separately. Where lines bordering the bars would overlap, the exact height and width of the bars are offset from each other so that each bar can be distinguished - it is intended as a visual guide, and the exact heights are therefore not reflective of the precise proportion of fins with an egg-spot in that position (see Table S3 for precise data).

When subsequent aggregations are initially adjacent, the central aggregations later reduce in cell density and disappear, with the flanking aggregations developing into separate egg-spots (Fig. 6A, white arrows). When initially distant, if a growing aggregation becomes close to another, a similar process of density reduction and cell disappearance occurs at the interface between the two aggregations (Fig. 6B, white arrows). Curiously, in one fin an aggregation started forming in the anterior spiny ray region, but similarly decreased in density and disappeared when a later-appearing aggregation in the typical first egg-spot position began maturing (Fig. 6D, white arrows). This shrinking of parts of aggregations in close proximity to others may indicate a form of competition between aggregations. Moreover, growing subsequent aggregations do not form steadily in all fins. After an initial appearance, aggregations may disappear for multiple days before forming again and maturing (Fig. 6C, grey arrows) or shift their position across a fin ray (Fig. 6D, green arrows).

To understand the impact of the more variable formation of subsequent egg-spots, we compared egg-spot positions between individuals in socialised cohorts at the end of the imaging series. At this stage most males show the adult phenotype: dense mature aggregations with a well-delimited boundary and outer transparent ring (n=10 out of 11) (Fig. S10). Of these, most males harboured three mature egg-spots (n= 9 out of 10) (Fig. S10), one individual of which showed an additional, later-appearing aggregation between the first and second mature egg-spots (Fig. S10K). The remaining male harboured two mature egg spots and a further small aggregation (n=1 out of 10) (Fig. S10B). We found that the position of the first egg-spot is the most consistent, always covering at least the region between the 4^th^ and 6^th^ fin rays (n= 10 out of 10), with most (n = 6 out of 10) extending more anteriorly, covering the region between the 3^rd^ and 6^th^ fin rays. One extended more posteriorly, covering the region between the 4^th^ and 7^th^ fin rays (Fig. 6E, Fig. S10, Table S3). The more posterior second and third egg-spots have more variability in their position at the end of our imaging series. The position of the second egg-spot was the most variable,with the most frequent position (n = 3 out of 10) being between fin rays 6^th^b and 8^th^b, followed by (n= 2 out of 10) 5^th^b to 7^th^b. All other positions for the second spot were unique to each individual (n = 6 out of 10) (Fig. 6F, Fig. S10, Table S3). The third spot position also had high variability. The most frequent positions covered the regions between 8^th^b and 9^th^b (n = 2 out of 9), 9^th^a and 10^th^ (n = 2 out of 9), and 8^th^b and 10^th^ (n = 2 out of 9) with the other positions unique to each individual (n = 3 out of 9) (Fig. 6G, Fig. S10, Table S3). All third egg-spots were on the distal posterior edge of the anal fin (Fig. S10).

This variability could be a result of the competitive interactions between growing aggregations; the earlier initiation and therefore greater size of the first aggregation may cause it to be less affected by these interactions, explaining the greater consistency in the position of the first egg-spot. Despite the variation in the precise position of the second and third egg-spots, the overall phenotype was very similar between males at the end of the imaging series (Fig. S10), indicating that similar outcomes can be achieved by a noisy formation process. In later adult stages, egg-spots continue to show apparently competitive interactions and can shift positions (Fig. S11) indicating that egg-spot patterns are dynamically changing throughout adulthood even after each egg-spot reaches a mature appearance.

## DISCUSSION

Elucidating the cellular and developmental processes underlying pigment pattern formation is essential to understand how variation can emerge. In this study, we were interested in characterising the development of the haplochromine cichlid egg-spots - a non-periodic male ornament harbouring high levels of intra and inter-specific variation. We focused on the egg-spots of *A. calliptera*, a cichlid fish part of the Malawi radiation that phenotypically resembles the radiation ancestor. We aimed to: 1) identify key cellular events contributing to egg-spot development and variation; 2) uncover when and how anal fin pigmentation development diverges between males and females; and finally 3) assess if external conditions, such as timing at which individuals became territorial, influences egg-spot development.

Overt egg-spot formation starts with the appearance of iridophores in the position of the future egg-spot (Fig. 7A-C). Egg-spot development then proceeds via the association of iridophores with xanthophore-like cells (Fig. 7D), a growing aggregation of associated cells (Fig. 7E) while xanthophore-like cells are lost from elsewhere in the fin (Fig. 7F), and finally the clearing of a transparent chromatophore-free ring (Fig. 7F). Contrary to our expectations, we found that both male and female juveniles exhibit the earliest stages of egg-spot development (small iridophore and xanthophore-like aggregations) (Fig. 7C-E)with anal fin patterns diverging only later between the sexes (Fig. 7F). Furthermore, the timing of egg-spot initialisation is plastic, with iridophore aggregations responding to the social conditions (Fig. 7C). Finally, there is variation between individuals in the development (Fig. 7G) and positions of the second and third spots.

**Fig. 7.**
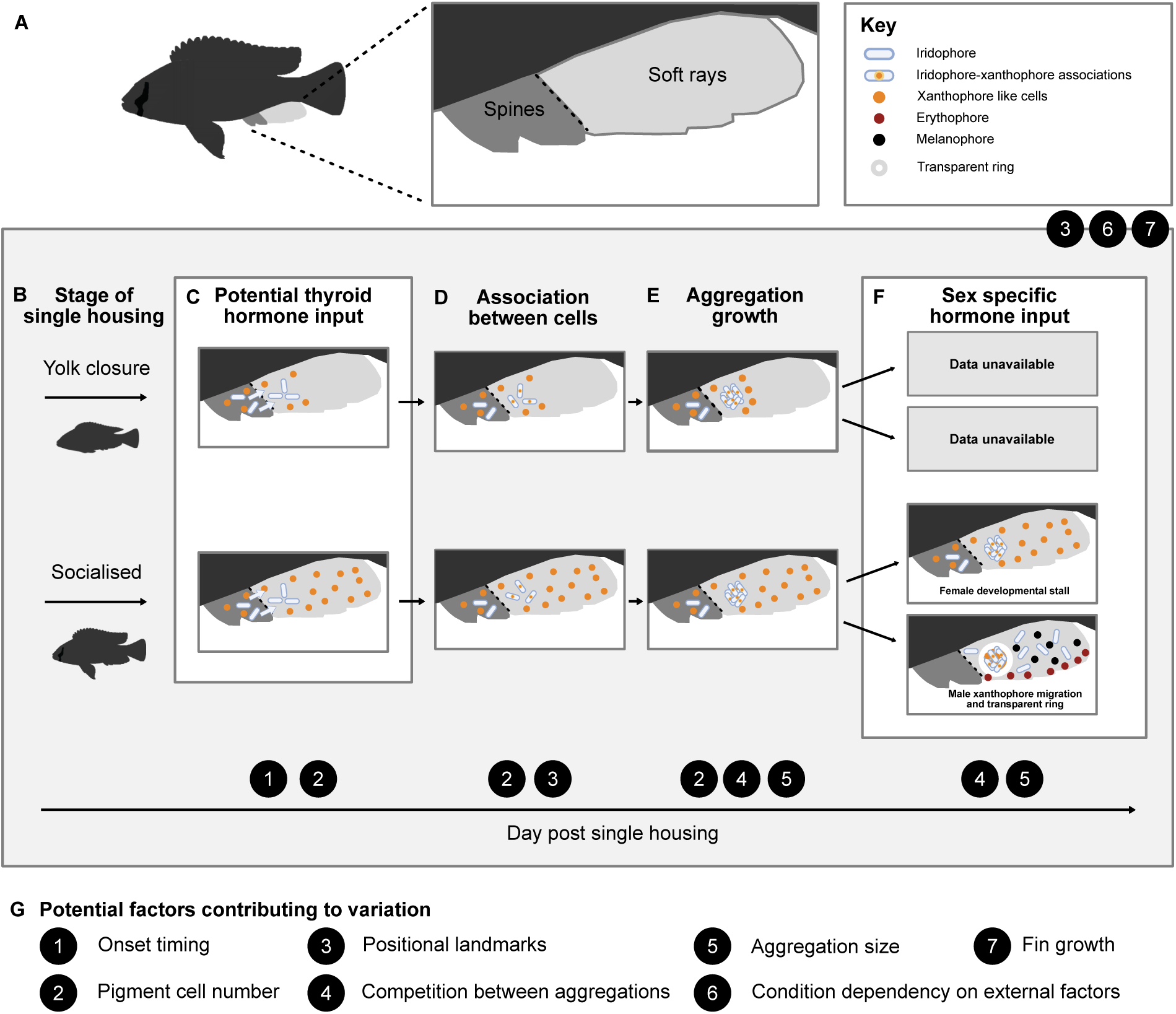
An overview of egg-spot development in *Astatotilapia calliptera*. (A) Representation of the spiny and soft fin ray domains in the anal fin. (B-F) Trajectory of egg-spot development in yolk closure and socialised cohorts, showing stages of egg-spot formation with hypothesised inputs. (G) Summary of hypothesised factors contributing to variation in adult phenotype and what stage of development they may act.

Cellular interactions play a significant role in the formation of periodic colour patterns in teleosts, but the role of positional information cues is hypothesised to be more important in the development of non-periodic patterns in order to define their position (Parichy, 2021). To evaluate the role of these mechanisms in egg-spot positioning, we deduce two scenarios with primary roles for cellular interactions or positional information, and consider cellular events during egg-spot formation for each scenario.

Iridophores first appear in the anterior hard-spined domain of the fin, likely migrating into the anal fin from the body as progenitors, but form aggregations in the soft-ray domain. Indeed, for the first egg-spot, we observed that an accumulation of iridophores occurs consistently between the fourth and fifth fin rays, posteriorly adjacent to the spiny-soft ray boundary in the anal fin. This suggests that the transition from a spiny-rayed to a soft-rayed tissue environment (Fig. 7A) plays a role in egg-spot patterning. The role of the soft-ray domain is supported by work in fin development in the Tanganyika cichlid, *Astatotilapia burtoni*: disruption to the signalling network that establishes the soft-ray domain causes concordant posterior shifts in both the spiny-soft ray boundary and the position of the egg-spots (Höch et al., 2021). We reason the soft-ray domain acts as a permissive or inductive region for iridophore aggregation following migration from the hard ray domain. In this scenario, egg-spots could emerge mostly due to cellular interactions between chromatophores, with iridophores initially aggregating and forming associations with xanthophore-like cells between the 4th and 5th fin rays solely because this is the first point of entry to this permissive/inductive environment. In an alternative scenario, within the soft-ray environment there could be multiple signalling centres inducing iridophores to aggregate in each spot location, one always present between the 4th and 5th fin ray and others in variable posterior positions (Fig. 7G-*2,3,5*). In either scenario, there may be tissue signals from fin rays as aggregations first begin to form between fin rays, and mechanisms to regulate the typically central proximal-distal positioning.

Both scenarios, cellular interactions and signalling centres, could explain the cellular events observed during egg-spot formation. The iridophore-xanthophore associations could result from short-range attraction between the two cell types or differentiation of iridophores into a novel cell type; in either scenario, this could be first triggered by entering the permissive zone or within close range of the first signalling centre. The aggregation growing in size and density could be due to migration of xanthophore-like cells and iridophores into the aggregations or local differentiation from unpigmented progenitors. In the latter case, the loss of xanthophores elsewhere in the fin may be due to cell death or loss of pigmentation, however we find no debris typical of dead pigment cells in our images. These mechanisms are compatible with both scenarios: cellular interactions can act at short- and long-range to promote local differentiation, migration, or cell death (Owen et al., 2020; Patterson and Parichy, 2013; Volkening and Sandstede, 2018; Yamaguchi et al., 2007). Notably, in zebrafish, iridophores attract xanthophores at short range (Patterson and Parichy, 2013). The same cellular behaviours can occur in response to a morphogen gradient from a signalling centre (Bier and De Robertis, 2015; Ninov et al., 2010).

Similarly, the clearing of a chromatophore-free transparent ring around the egg-spot in males could be due to repulsion, induced cell death or depigmentation in response to cellular interactions or signalling centres. In zebrafish, both iridophores and xanthophores repel melanophores at long range while attracting each other at short range to separate light interstripes from dark stripes (Frohnhöfer et al., 2013). Similar interactions in *A. calliptera* fins could explain the unpigmented region around the maturing egg-spot, the transparent ring. Alternatively, a diffusible signal from the signalling centre or the aggregation itself could be decoded by concentration threshold to induce cell death or repulsion in the ring but not beyond it. Both scenarios require an explanation for the change in aggregation behaviour from growth to ring clearing. It may be that as the aggregation reaches a certain density or cell number, the signals/interactions from the aggregation reach a critical threshold or even change in nature. Consideration of these various events shows that either scenario of self-organising cellular interactions or tissue landmarks patterning mechanisms could underlie the development of egg-spot patterns, and in fact demonstrates that each scenario would likely involve both mechanisms. Analyses of pigment cell mutants in *A. calliptera* will be required to understand the relative contribution of each mechanism and the specific contribution of each cell type for egg-spot formation.

That said, the variability in the formation and final position of the second and third egg-spots supports the predominantly cellular interactions scenario. Most notably, aggregations shifting position across fin rays while forming is more parsimonious with dynamic self-organisation than positioning being controlled by a predefined signalling centre. The apparently competitive interactions between developing aggregations are a further indication of the role of interactions between pigment cells. Rather than subsequent aggregations initiating in response to multiple signalling centres, iridophore-xanthophore associations can initially form in anterior-posterior locations that can differ between individuals, and competitive inter-aggregation interactions determine which aggregations in each fin develop into egg-spots (Fig. 7G*-4*). The variation observed in this process of subsequent spot formation may contribute to the adult intra-specific variation in *A calliptera* egg-spot, including spot number and precise position. Moreover, the overall resulting patterns are recognisably similar despite this variability during formation and in precise final position, which also indicates a self-organising system driven by cellular interactions rather than a determined pre-pattern.

The question that arises is how this self-organising process leads to variation in egg-spot position and number, unlike zebrafish colouration self-organisation which gives rise to highly reproducible and precise periodic stripes. Greater noise, or susceptibility to noise, in egg-spot formation could explain this (Maini et al., 2012), such as inter-individual variation in external conditions, hormonal levels, system responsiveness to these two factors, or pigment cell interactivity. There also remains the question of understanding how this variability in egg-spot formation manifests in the development of later egg-spots after the third forming egg-spot. Growth may be a factor: as the area of the fin increases this may create enough space between existing egg-spots for new aggregations to develop without competition from their larger neighbours (Fig. 7G*-6, 7*). Comprehensive imaging long-term of later egg-spot development is necessary to address this question.

Like many other aspects of dominant male colouration in cichlids, egg-spots are most strongly expressed when a male is dominant. In our study, we found that the timing of egg-spot initiation is plastic and dependent on the social environment with iridophores aggregating in the position of the first egg-spot upon single housing. When isolated, the fish effectively becomes dominant in its single-housing territory, which suggests that the onset of egg-spot development depends on social status. This timing plasticity may be a mechanism by which some smaller fish sometimes bear higher numbers of egg-spots than larger fish in *A. calliptera:* if an individual becomes territorial earlier, they would start adding egg-spots at earlier stages and smaller standard lengths (Fig. 7G*-1*). Orange pigmentation traits are often sexually-selected honest indicators of condition because the orange colour of xanthophores depends on dietary uptake of carotenoids (Leclercq et al., 2010; Olson and Owens, 1998). Therefore it is surprising that underlying plasticity in egg spot development timing, iridophores are the condition-dependent cells that respond to the social environment cue.

There is likely hormonal mediation between sensing social conditions and the aggregation of iridophores, as hormones often mediate developmental plasticity (Brakefield et al., 1998; Ledón-Rettig and Ragsdale, 2021; Moczek and Nijhout, 2002). A likely candidate is thyroid hormone, as in zebrafish thyroid hormone promotes iridophore maturation as well as limiting population expansion of melanophores and promoting accumulation of carotenoid pigment in xanthophores (McMenamin et al., 2014; Saunders et al., 2019). Further, in clownfish thyroid hormone mediates between environmental conditions and speed of formation of iridophore-based white bars (Salis et al., 2021). A similar mechanism could regulate iridophore aggregation behaviour at the site of the future egg spots.

Contrary to our expectations, we observed social isolation-induced iridophore aggregations in both males and females, showing this event is a developmental program of fin patterning common to both sexes. We therefore speculate that a sex-independent increase in thyroid hormone regulates iridophore aggregation behaviour (Fig. 7C). Differences between males and females emerge later in development when the iridophore-xanthophore aggregations grow in size, xanthophore-like cell distributions change and the transparent rings form in males. A sex-independent capacity to initiate egg-spot development in response to social dominance but a sex-limited capacity to develop mature egg-spots explains why immature egg-spots sometimes occur on the anal fin of dominant females (Heule and Salzburger, 2011). These late developmental differences between sexes are likely modulated by sex-specific hormones (Fig. 7F). For example, in the brown anoles lizard, a female-specific pigmentation polymorphism is regulated by co-expression of the *estrogen receptor-1* gene with *ccdc170* (Feiner et al., 2022). Similarly, in cichlids, up-regulation of sex-steroids and gonadotropins in males are associated with a rise to dominant social status and development of sexual dimorphism (Maruska et al., 2011; Maruska et al., 2022; Moore et al., 2022).

Taken together, our data indicates that egg-spots are a promising system to investigate teleost colour patterning development and evolution. It appears likely that egg-spot development largely self-organises by cellular interactions, and is initiated by iridophores, similar to the well-studied formation of zebrafish stripes. With previously undescribed cellular morphologies and behaviours including variably contracting xanthophore-like cells, iridophore-xanthophore associations, and transparent ring clearance, egg-spot formation lends further evidence to the suggestion that pigment cells and their interactions may be evolutionarily labile (Kelley et al., 2013; McCluskey et al., 2021; Singh and Nüsslein-Volhard, 2015). Moreover, the fact that egg-spot formation can start regardless of the stage of background xanthophore-based pigmentation development contradicts a key tenet of zebrafish stripe formation in which contributions from all aspects of pigmentation are required for the pattern to emerge (Patterson and Parichy, 2019). In addition to self-organising inter-cellular interactions, we propose multiple additional inputs to regulate the position, timing, and maturation of the egg-spot pattern: an inductive/permissive tissue domain to bound initial iridophore positioning, sensing and hormonal mediation of social conditions influencing timing of pattern initiation, and a sex-limited hormone enabling egg-spot maturation only in dominant males. Therefore, egg-spots provide an exciting model for investigating how environmental and conspecific cues affect pigment pattern formation at the cellular and developmental level. Furthermore, *A. calliptera* represents the ancestral bauplan for egg-spots. As such our findings provide a baseline for informed comparisons across the Malawi radiation which harbours an incredible extent of inter- and intra-specific variation in egg-spots. Addressing the genetic and developmental underpinnings of such variation will yield insights into the genes, signals, and cellular interactions that underlie the mechanisms patterning non-periodic male pigmentation ornaments and their evolution.

## MATERIALS AND METHODS

### Fish housing

*Astatotilapia calliptera* stocks were kept under constant conditions (25 ± 1°C, 12 h dark/light cycle, pH 8) in 220 l tanks. Fish were fed twice a day with cichlid flakes and pellets (Vitalis). Tank environment was enriched with plastic plants, hiding tubes, and coral sand substrate. Embryos were extracted from mouthbrooders and raised in tumblers until free-swimming juveniles.

### Experimental design

Three imaging series were completed, starting at three different developmental stages (Fig. 2). For each developmental stage, two replicates were performed, one with shallow individuals *A. calliptera* ‘masoko’ and one with deep individuals. Individuals were kept in typical social groups of single or multiple clutches until the day imaging started. At the onset of each imaging series individuals were housed individually to keep track of each individual through time. *<u>Embryo cohort</u>*: Embryos were kept in separate egg tumblers and imaged daily, from when the anal fin condensation becomes visible (10 days post fertilisation, dpf) until the transition to the free-swimming juvenile stage, marked by yolk closure where the skin closes over the yolk. *<u>Yolk closure cohort</u>*: Juveniles were housed individually upon reaching the free-swimming stage, at the point of yolk closure, and imaged every 3 days until 12 days after yolk closure and every 5 days thereafter. Juveniles were housed in cylinders (14 cm diameter) constructed of plastic mesh allowing for water flow. All cylinders were placed in the same aquarium (220 l) so that the water conditions and temperature were the same for all the individuals in each imaging series. For the first 15 days of imaging, a nylon net was placed within the cylinder, using material from tights and weighed down with sand, to prevent the fish escaping the cylinders due to their small size. Once larger, the net was removed and plastic plants were added to each cylinder as enrichment. *<u>Socialised cohort</u>*: Juveniles were kept in a social group until 6 weeks post yolk closure for deep individuals and until 10 weeks post yolk closure for shallow individuals. These individuals were then individually housed in the mesh cylinders described above and imaged every 5 days. Three to five days was the maximum imaging frequency for juveniles because daily imaging affected survival. The sample sizes for each cohort were as follows: 1) Embryo cohorts: shallow n = 8, deep n = 8; 2) Yolk closure cohort: shallow n = 8 (5 male, 3 female), deep n = 6 (3 male, 3 female); Socialised cohort: shallow n = 12 (5 males, 7 females); Deep n = 9 (5 males, 4 females from two clutches).

Yolk closure cohorts and the shallow socialised cohort imaging series spanned 50 days; the deep socialised cohort spanned 150 days. The shallow socialised cohort was conducted first as a pilot imaging series, with individuals selected for visibly initiated egg-spot development (existing aggregations) to maximise the likelihood of capturing the completion of egg-spot development. Following assessment of the images, we planned the timing of other cohorts and deep/shallow replicates with the aim of capturing all stages of egg-spot development from fin condensation to mature spots. Length of imaging series for each cohort was limited by survival rates and project licence conditions. It was not possible to continue imaging embryo cohorts after yolk closure, because repeated anaesthesia as juveniles after daily anaesthesia as embryos caused low survival rates. The project licence limited juvenile series to 50 days initially, but was amended to 150 days for the deep socialised cohort in order to capture as much of egg-spot development within the same individuals as possible.

### Sex genotyping

DNA was extracted from fin clips stored in 100% ethanol for <2 weeks using the PCR biosystems rapid extract lysis kit or Quick-DNA^TM^ Miniprep Plus kit by Zymo research. *Astatotilapia calliptera* “kisiba/masoko” has three XY sex-determining systems (Munby et al., 2021). A multiplex PCR assay was developed with three different primer sets to genotype the three different sex genotypes (Table S1). Two primer sets target a *gsdf* duplication and TE insertion on chromosome 7 and the other set a TE insertion on chromosome 19. PCR and subsequent gels for visualisation were run with negative and positive controls for each of the sex determination systems present in the species (Munby et al., 2021). The master mix contained Platinum™ II Taq Hot-Start DNA Polymerase (Invitrogen^TM^ (Thermo Fisher Catalog number: 14966001) The PCR program was 94°C for 2 min, 30-35 cycles: 94°C for 15 seconds; 60°C for 15 seconds; 68°C for 15 seconds; Final extension: 68°C for 5 minutes. PCR products were visualised on a 2% agarose gel with ethidium bromide at 100V for 30 minutes. All heterozygous samples should have the reference genome band (∼400bp). Presence of insertion is indicated if a smaller band is visible in both the chromosome 7 and 19 TE insertions. *Gsdf* duplication presence is indicated by smaller (207bp) and larger (614bp) bands than the reference.

### Anaesthesia procedure

Fish were immersed in epinephrine (Sigma-Aldrich (Merck) Product number E4250) (1 mg/ml) to contract pigment cells alongside 0.01% Tricaine (Ethyl 3-aminobenzoate methanesulfonate, Sigma-Aldrich (Merck) Product number E10521) for 5 minutes, then anaesthetised in 0.02% tricaine until immobilised (approximately 1 minute). Fish were imaged for <30 minutes in a petri dish in 0.01% tricaine to maintain anaesthetisation then allowed to recover in tank water. Exposure to epinephrine for 5 minutes is not sufficient to contract all pigment cells, but longer exposure greatly increases the chance of death, so fish would be unlikely to survive repeated exposure.

### Imaging and image processing

Images were taken with a Leica M205 FCA stereoscope with DFC7000T camera. Standard length images of the whole body were taken at 1X, with multiple tiles taken if the specimen was too large to be contained within a single frame. Anal fin images were taken at 6.3X magnification at multiple focal planes and multiple tiles. All fins were imaged under both darkfield reflected and brightfield transmitted light. Each tile of focal planes was merged using the batch process function in Helicon Focus 8, with Method B, radius 8, smoothing 4. Then, the tiles for a single individual were stitched together using the panorama function in Affinity Photo version 1.10.5.

### Event scoring and standard length measurement

Events during fin development were defined as: *<u>Xanthophore-like cell appearance</u>*: any xanthophore-like cells visible in the fin, including both the contracted orange circle (responded to epinephrine treatment) and the non-contracted appearance of a dispersed orange hue (no epinephrine treatment or non-responsive) (Figure S1). *<u>Iridophore appearance</u>*: any iridophore visible in the fin, identified as grey lines in images taken under transmitted light and confirmed as reflective in images taken under reflected light. *<u>Iridophore appearance at specific positions</u> <u>in the fin</u>*: any iridophore visible between the designated numbered fin rays. *<u>Xanthophore-</u> <u>iridophore association</u>*: when xanthophores directly overlap with densely-accumulated iridophores. *<u>Clear outer transparent ring</u>*: complete ring without chromatophores surrounding the eggspot.

For each individual, the recorded event timing reflects the earliest image showing the defined characteristic of each event. Standard length was measured from the tip of the snout to the caudal peduncle in whole-body stereomicroscope images or photographs, using the straight line tool in Fiji (Schindelin et al., 2012). Event timings relative to days and standard lengths were plotted in R using ggplot2 (Wickham, 2016). Differences in event timings between cohorts were tested in R using the Kruskal-Wallis test when only two groups were being compared, and pairwise Wilcox with Bonferroni adjustment when multiple groups were compared (Table S2). Event scoring data is available in Supplementary File 1 and also on GitHub (https://github.com/Santos-cichlids/Developmental-plasticity-and-variability-in-egg-spot-formation-in-Acalliptera).

### Image segmentation

Image segmentation for quantification of melanophores and xanthophores was performed using Affinity Photo, ilastik and Fiji. Melanophore coverage was quantified across the whole fin in stages before egg-spots began developing; for each image the background was removed using the selection brush tool in Affinity Photo. Cell segmentation was performed using the machine learning image segmentation tool ilastik (Berg et al., 2019). One image from each imaging day was used for the training dataset using the pixel classification tool. Feature selection parameters colour/intensity, edge and texture were all given a sigma of 10. Iterative learning cycles were performed until the cell type of interest was appropriately classified. To compensate for background colouration changes between imaging sessions, images were batch processed by imaging session. Outputs were exported as simple segmentation tiff files and reimported to Fiji for quantification of chromatophore coverage and number using the Analyse Particle function.

Quantification of xanthophore-like cell distribution was performed in a region outside of the developing egg-spots to avoid noise from other chromatophore types. Only a defined region of the fin was measured as a proxy for xanthophore-like cell distribution across the anal fin: a 0.5mm^2^ square over the first segment of the eighth fin ray was extracted from each image using Fiji.The outside of the image was cleared and the remaining 0.5mm^2^ image exported. For both xanthophore-like cell morphologies, the extracted images were imported into ilastik. Two iterations of the pixel classification tool in ilastik on each xanthophore-like cell type were performed, with one individual from each imaging session selected for the training set as described above. Cell counts and area measurements were performed in Fiji as described above, and the sum of the area of both xanthophore-like cells was used as the total area coverage within the sampled region. To analyse the effect of sex and time since single housing on xanthophore coverage we used Pearson’s correlation to measure the strength and direction of xanthophore-like cell area coverage over time. For further details see our GitHub page (https://github.com/Santos-cichlids/Developmental-plasticity-and-variability-in-egg-spot-formation-in-Acalliptera).

## Acknowledgements

We thank the Animal Technicians and NACWO at the UBS Cichlid fish facility in Madingley (Cambridge) for the help in rearing fish and Members of the Morphological Evolution Lab for technical support and discussion. We would also like to thank Walter Salzburger, David Parichy and Dylan Huang for reading and providing very constructive feedback on the manuscript.

## Ethics

All animals were handled in strict accordance with the protocols listed in the Home Office project licence PCA5E9695.

## Data accessibility

The datasets supporting this article have been uploaded as part of the electronic supplementary material. Raw image data can also be found at Dryad repository.

## Author’s contributions

AH, BC, MES contributed to study conceptualisation, funding acquisition, investigation, writing, review and editing of the manuscript. BF and JE contributed to data acquisition. All authors read and approved the final version of the manuscript.

