## Supplementary Figures and Tables for "Developmental plasticity and variability in the formation of egg-spots, a pigmentation ornament in the cichlid *Astatotilapia calliptera*"

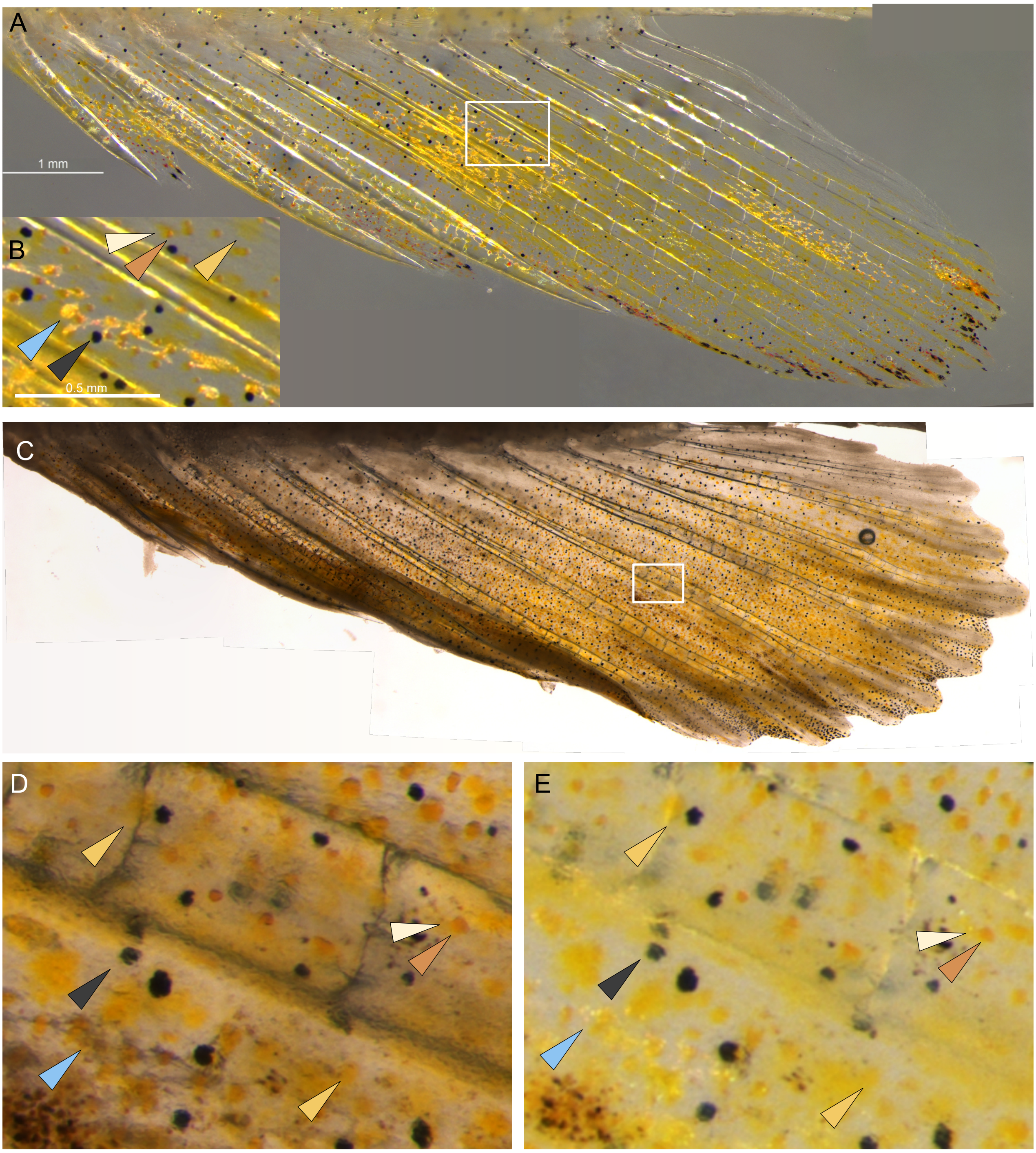


**Fig. S1. Different xanthophore-like morphologies under different lighting conditions and epinephrine treatments.** (A-E) Fins subjected to longer or higher epinephrine treatments to test the responsiveness of xanthophore contraction to epinephrine. In both, uncontracted xanthophores are still visible (light yellow arrow) as well as contracted melanophores (black arrow) contracting xanthophore-like cells (orange arrow) and iridophores (grey arrow). For the contracting xanthophore-like cells, relief illumination by Rottermann’s Contrast technology makes the surrounding ‘halo‘ of lighter pigment (pale yellow/white arrow) more visible. (A-B) A fin treated with normal levels of epinephrine 1mg/ml, imaged after several hours under relief illumination with additional reflected light. (C-D) A fin treated with high levels of epinephrine (2.5mg/ml) imaged after 30 minutes under white transmitted lighting (C-D) and relief illumination with additional reflected light (E).

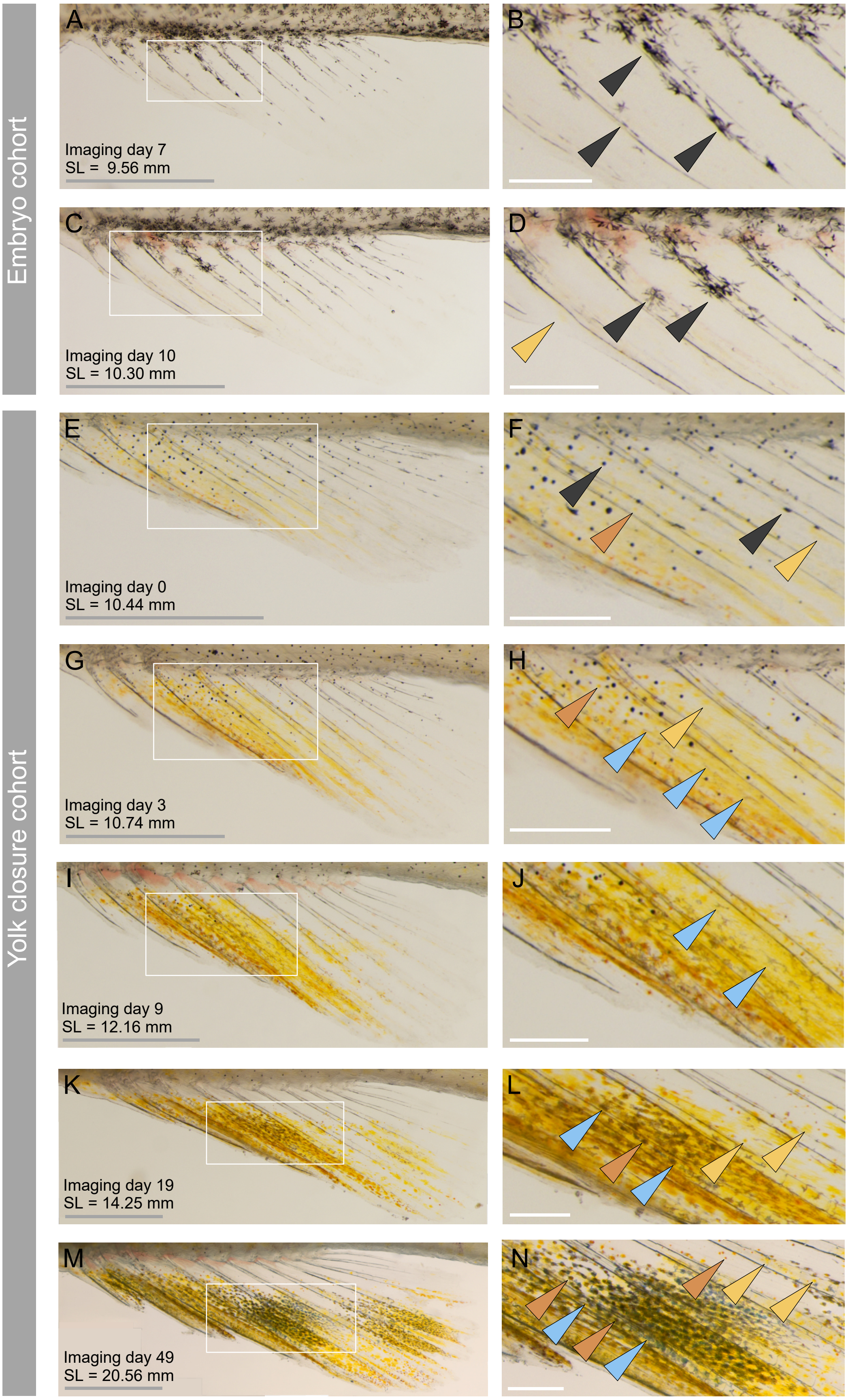


**Fig. S2. Description of early anal fin pigmentation ontogeny leading up to the initial stages of egg-spot development in *A. calliptera* shallow individuals.** (A-N) Images with transmitted light of developing fins showing different stages of spot initialisation in embryos (A-D) and yolk closure cohorts (E-N). Events shown are melanophores present in embryo fin (A-B), xanthophore-like cells present in embryo fin (C-D), xanthophore-like cells present in yolk-closure fin (E-F), iridophores first present in fin (G-H) iridophores first present in position of first spot, i.e. between fourth and fifth fin ray (I-J*)*, iridophores appear stellate and associate with xanthophore-like cells (K-L), egg-spot aggregation at end of 50 day imaging isolation period (M-N). Arrows indicate melanophores (black), contracting xanthophore-like cells (orange), uncontracting xanthophore-like cells (yellow), and iridophores (blue) . Grey scale bars 1 mm, white scale bars 250 µm. Embryo fins are not treated with epinephrine, early juvenile fins are treated with epinephrine causing the pigments in the chromatophores to contract.

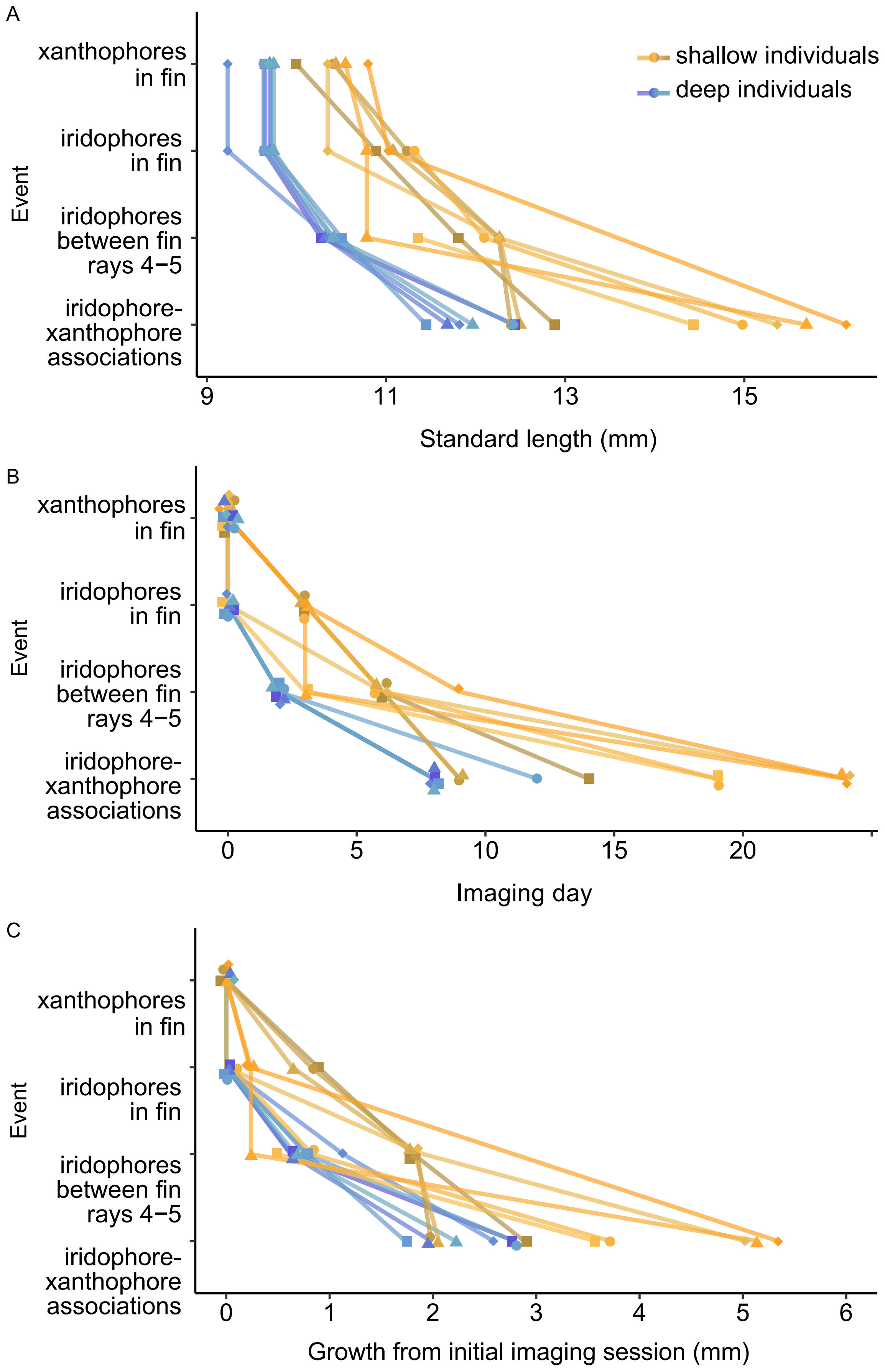


48 **Fig. S3. Order of events during egg-spot initalisation is consistent between individuals**

49 **and populations.** (A-C) Graphs indicating the developmental trajectory of early pigmentation

events in deep and shallow yolk closure cohorts by individual. For the first imaging session in which the event was observed, each event is plotted against standard length (A), days since first imaging session (B), and increase in standard length from first imaging session (C). Differences between shallow and deep cohorts when measured in SL (A) are resolved when normalising for starting SL (C*)* similarly to measuring by days (B). The timing of xanthophore appearance and iridophore appearance in the fin of yolk closure cohorts is more closely aligned between individuals when measured as days after the initial imaging session (B) or growth since initial imaging session (C) instead of by standard length (A). Note this differing alignment is an artefact that reflects the fact that for most individuals these events were recorded on the initial imaging session, likely representing earlier occurrence in embryo stages. For events occurring after yolk closure, all three measurements of developmental progression (A-C) are similarly good predictors of event timing within the yolk closure cohorts.

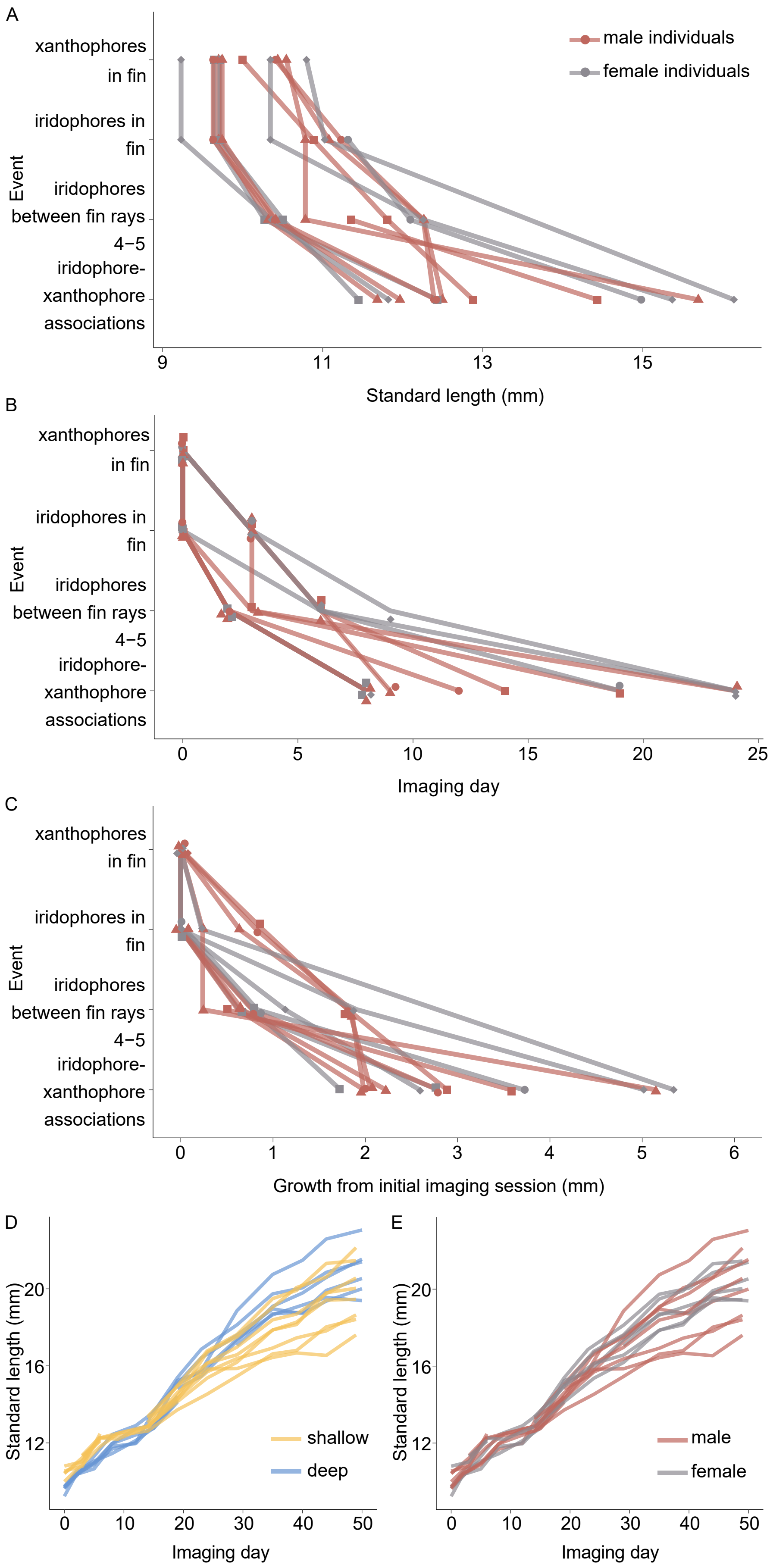

A
male individuals
female individuals
xanthophores in fin
iridophores in fin
iridophores between fin rays 4–5
iridophore-xanthophore associations
Event
9
11
13
15
Standard length (mm)
B
xanthophores in fin
iridophores in fin
iridophores between fin rays 4–5
iridophore-xanthophore associations
Event
0
5
10
15
20
25
Imaging day
C
xanthophores in fin
iridophores in fin
iridophores between fin rays 4–5
iridophore-xanthophore associations
Event
0
1
2
3
4
5
6
Growth from initial imaging session (mm)
D
Standard length (mm)
12
16
20
shallow
deep
0
10
20
30
40
50
Imaging day
E
Standard length (mm)
12
16
20
male
female
0
10
20
30
40
50
Imaging day

**Fig. S4. Growth and order of events during egg-spot initalisation is consistent between sexes.** (A-C) Graphs indicating the developmental trajectory of early pigmentation events in deep and shallow yolk closure cohorts by sex. For the first imaging session in which the event was observed, each event is plotted against standard length (A*)*, days since first imaging session (B), and increase in standard length from first imaging session *(C).* In trajectory graphs, the result of male and female individuals are intermingled. *(*D-E) SL graphs indicating standard length of each individual in the yolk closure cohorts against days since first imaging session, coloured by morph (D) and sex (E*)*. In SL growth graphs, results are intermingled for shallow and deep individuals, likewise male and female individuals.

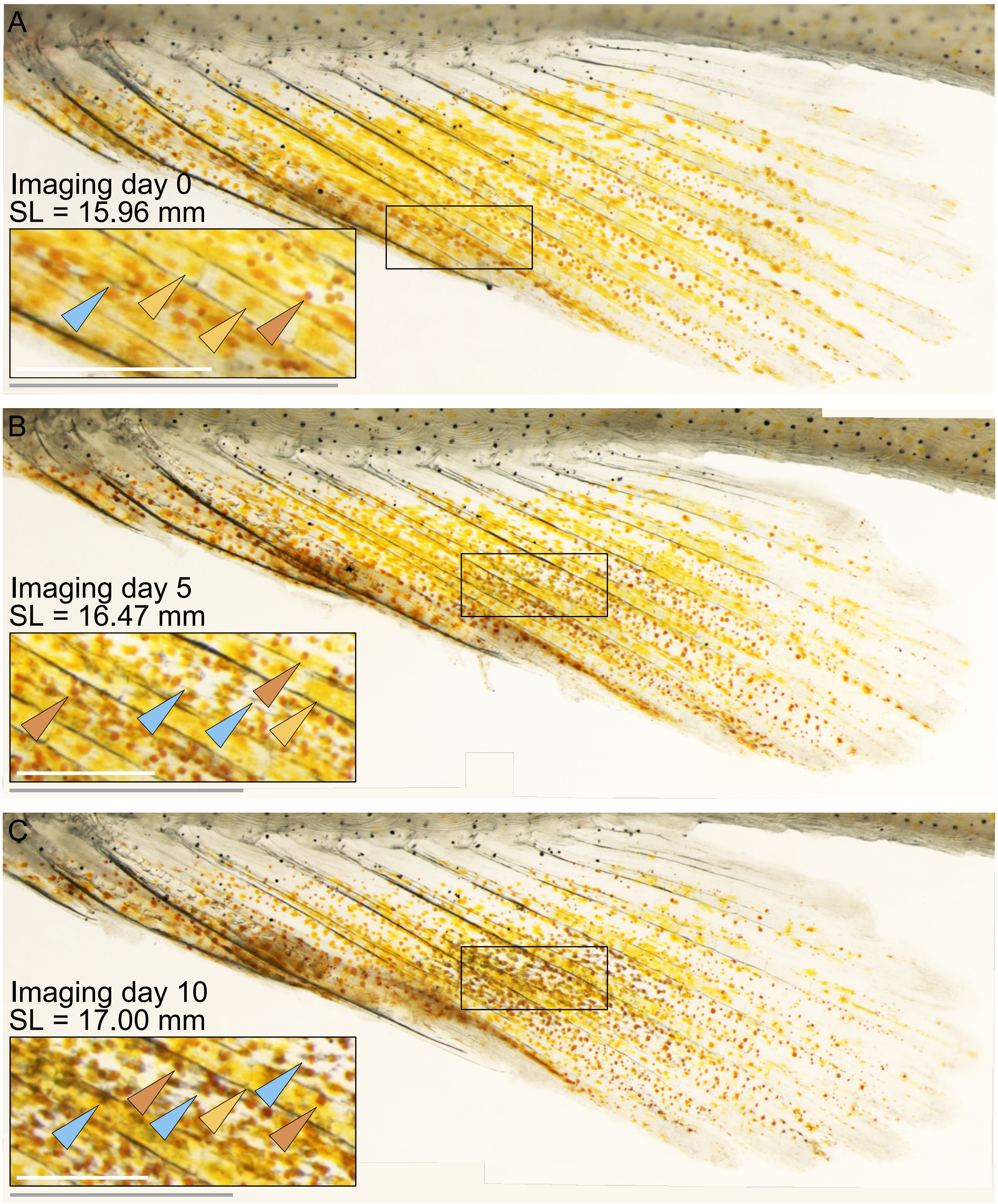


**Fig. S5. Initial stages of egg-spot development in socialised cohorts are consistent with yolk-closure cohorts .** (A-C) Images with transmitted light of developing fins showing egg-spot initialisation in a deep morph individual from the socialised cohort: xanthophores and iridophores first present in fin (A), iridophores first present in position of first spot, i.e. between the fourth and fifth fin ray (B), and appearance of iridophore-xanthophore associations (C).

Arrows indicate contracting xanthophore-like cells (orange), uncontracting xanthophore-like cells (yellow), and iridophores (blue). Grey scale bars 1 mm, white scale bars 250 μm.

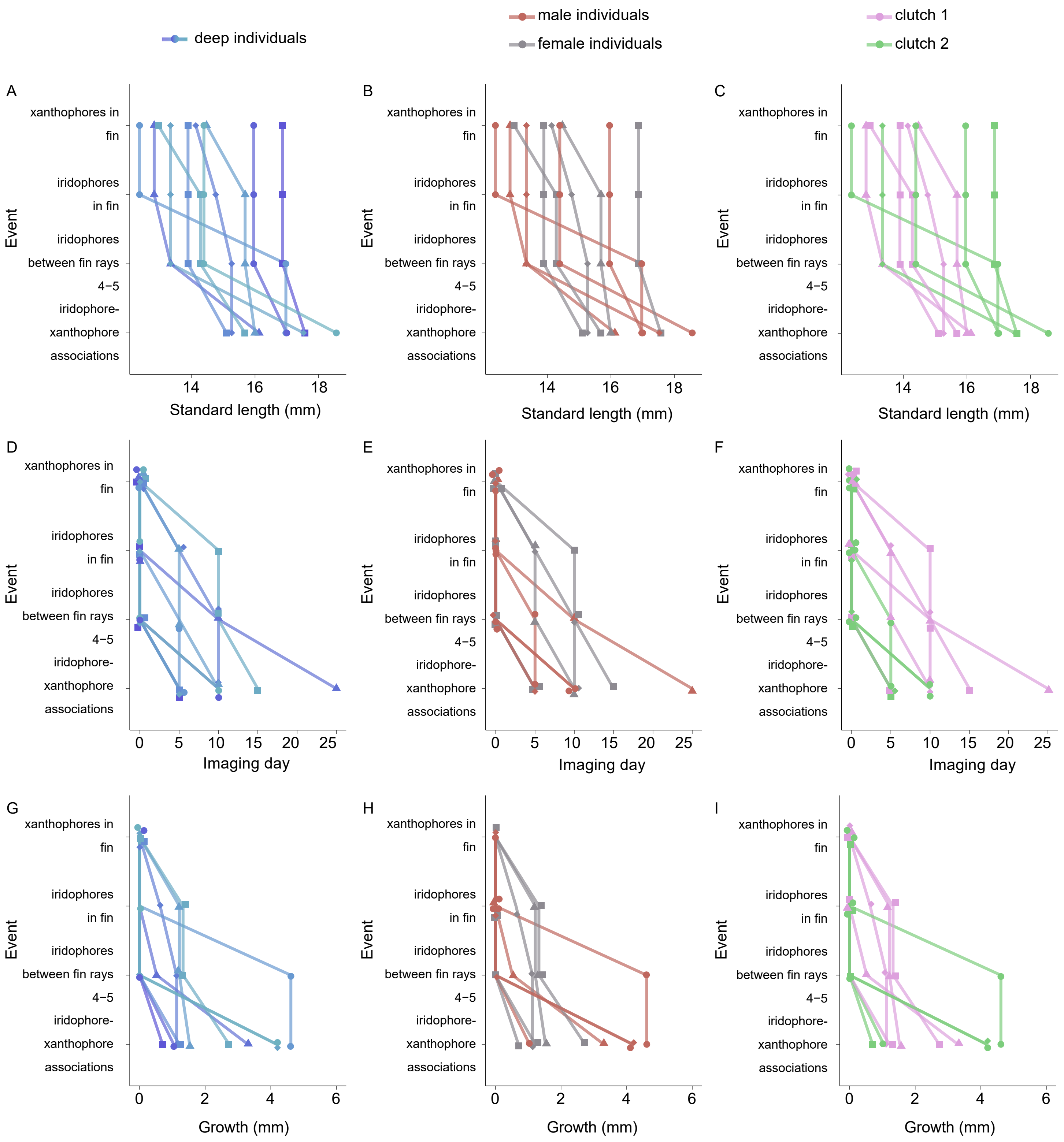


**Fig. S6. Order of events during egg-spot initalisation is consistent between and within cohorts.** (A-I) Graphs indicating the developmental trajectory of early pigmentation events in deep socialised cohorts. For the first imaging session in which the event was observed, each event is plotted against standard length (A-C), days since first imaging session (D-F), and increase in standard length from first imaging session (G-I). Colouring by individual (A,D,G) shows event order is consistent between individuals within the deep socialised cohort and between the socialised and yolk cohorts in Fig 3. Colouring by sex (B,E,H) shows the trajectories for male and female individuals are intermingled for all developmental progression measurements. Colouring by clutch (C,F,I) shows the trajectories are intermingled for developmental progression measurements standard length and growth but not by day, by which some clutch 1 individuals exhibit a 5-15 day delay in iridophore events.

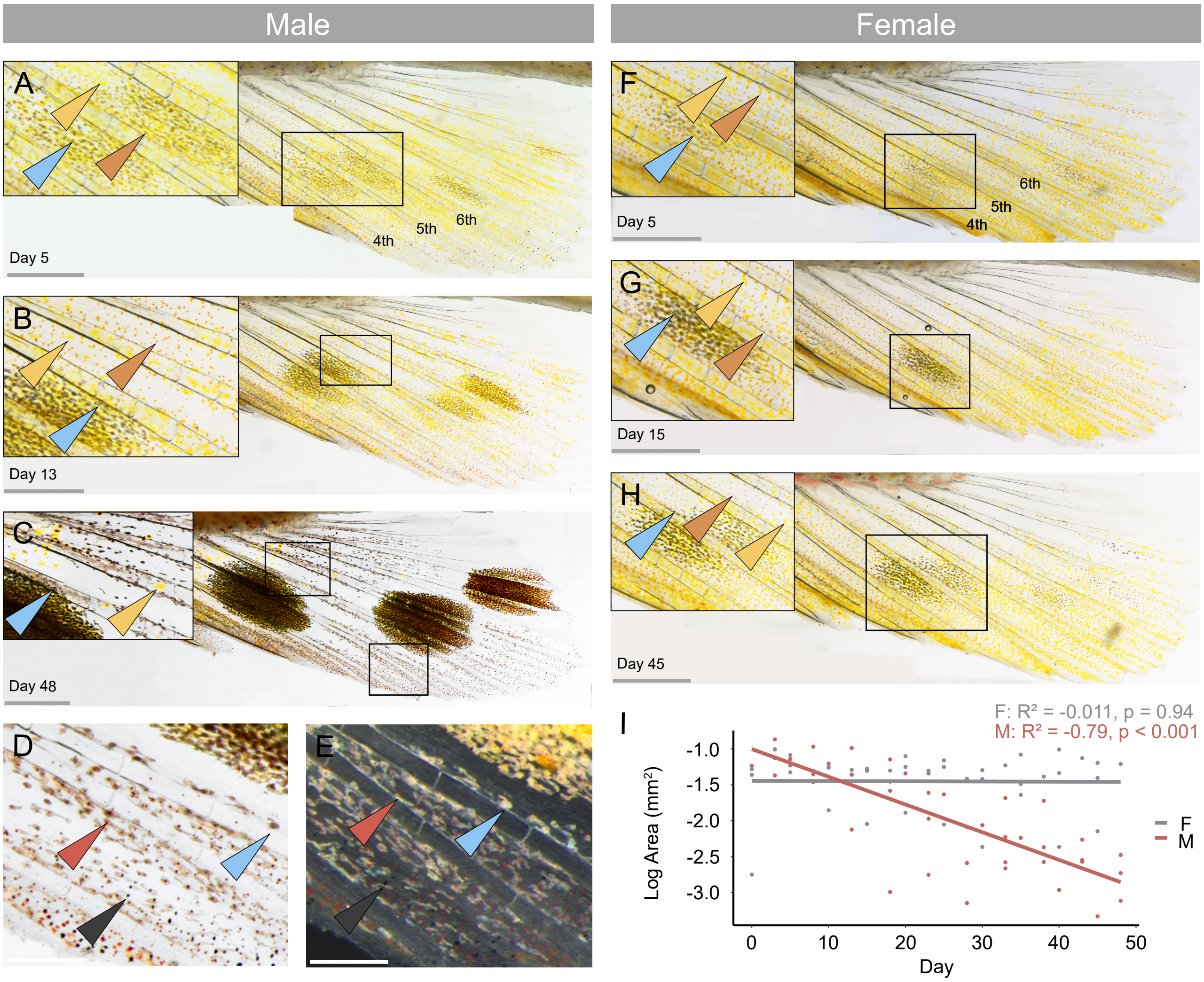


**Fig. S7. Divergence in egg-spot development between sexes in A. *calliptera* "masoko/kisiba" shallow individuals showing consistency in later-egg spot events between morphs** Images indicate shared development of and point of divergence in juvenile (A-C) male and (F-H) female A.calliptera *kisiba* shallow anal fins, similarly to deep morph individuals in figure 4. Black boxes highlight the first egg-spot aggregation region between soft rays 4-6. White arrows: stellate associated iridophores; blue arrows: elongate unassociated iridophores; yellow arrows: light variably contracting xanthophores; orange arrows: dark contracting xanthophores; black arrows: melanophores; red arrows: erythrophores. Elongate unassociate iridophores, erythrophores and melanophores (E-D) are found in mature male individuals (C) outside of the egg-spot region. (I) Area coverage of both xanthophore morphologies was measured in a $0.5mm^2$ square region centred on the first segment of the eighth fin ray. An inverse correlation was found between xanthophore coverage and Day in males (pearson's correlation R2 = -0.79, $p < 0.001$) but not in females (R2 = -0.011, $p = 0.94$)(See GitHub).

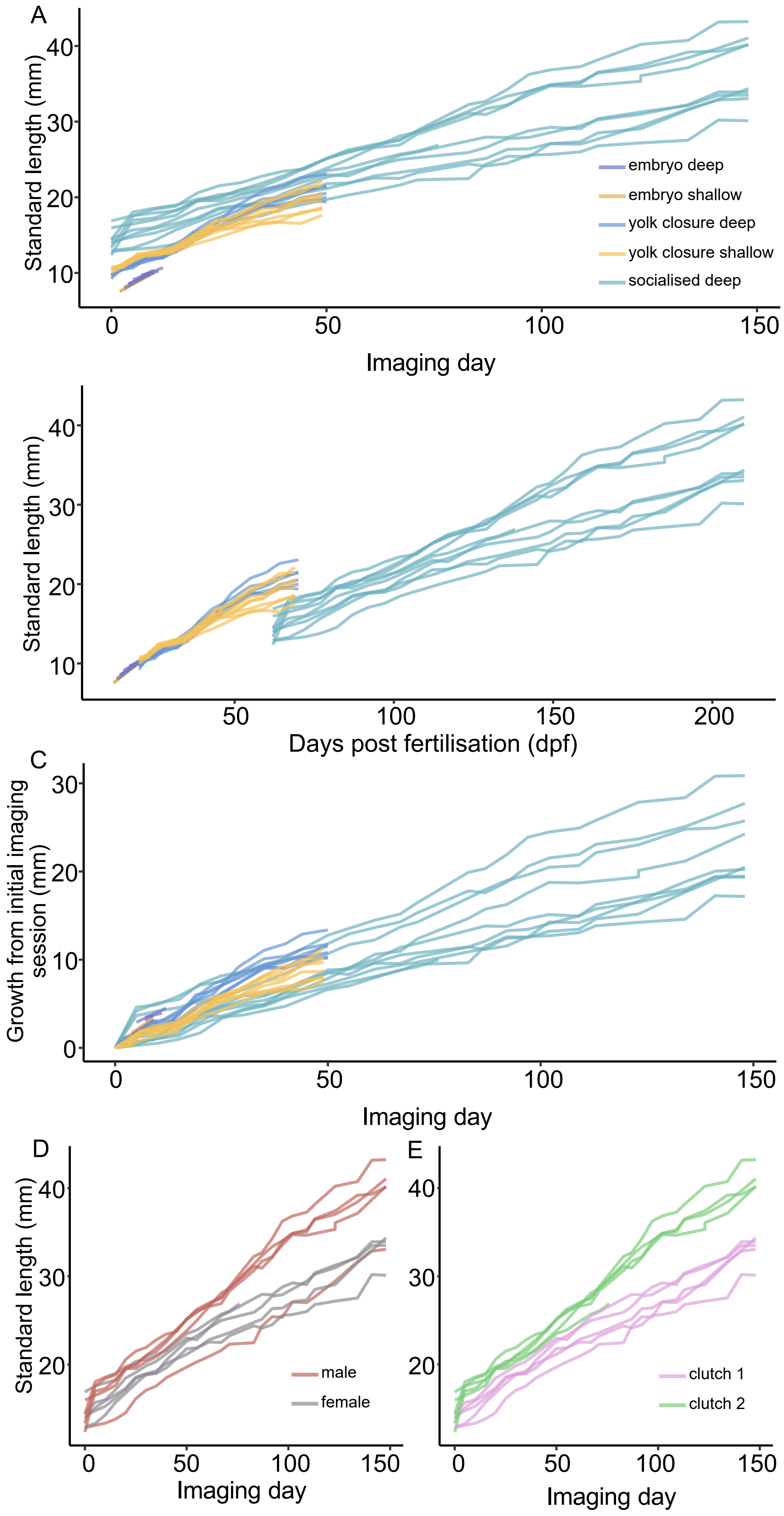
A
Standard length (mm)
Imaging day
embryo deep
embryo shallow
yolk closure deep
yolk closure shallow
socialised deep
Standard length (mm)
Days post fertilisation (dpf)
C
Growth from initial imaging session (mm)
Imaging day
D
Standard length (mm)
Imaging day
male
female
E
Imaging day
clutch 1
clutch 2

**Fig. S8. Growth rates are consistent between and within cohorts**. Standard length is a descriptor of developmental stage, which can be reliably applied across cohorts as growth rates are comparable. Moreover, standard length more accurately reflects developmental stage than dpf as starting and ending sizes are not aligned between yolk and socialised cohorts, and in fish developmental progress depends on size and growth rate.(A) Standard length of each individual, coloured by cohort, against days since first imaging session. Length measurements overlap between morphs and individuals for each cohort. For adult cohorts (yolk-closure and socialised) there is a small increase in standard length variation between individuals as time progresses. (B) Standard length of each individual, coloured by cohort, against dpf. Individuals at the start of the socialised cohort are smaller than the end of the yolk closure cohort, likely because growth rates are slower during time spent in a social group. (C) Increase in standard length from initial imaging session individual, coloured by cohort, against days since first imaging session. Growth measurements overlap between all morphs and cohorts. Standard length measurements were not obtained for the shallow socialised cohort. (D-E) Standard length of each individual in the deep socialised cohort against days since the first imaging session, coloured by sex (D) and clutch (E). The deep socialised cohort consisted of two clutches. Clutch differences, not sex, explain the divergent growth rates within the socialised deep cohort.

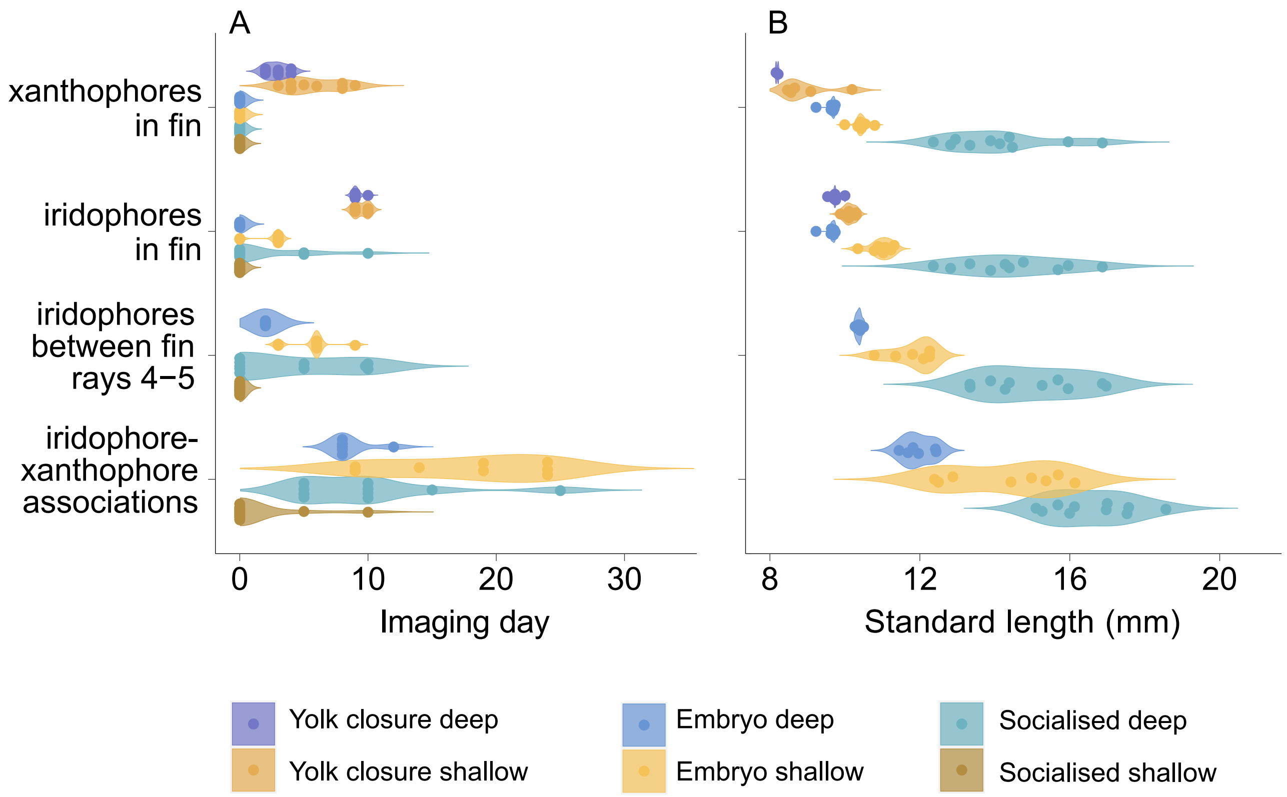


**Fig. S9. Timing of egg-spot initialisation events.** Violin plots showing event timing by cohort and morph, against contrasting measures of days in single housing (A) and standard length (B). For the first imaging session in which the event was observed, each event is plotted against days since first imaging session (Day) (A), and standard length (SL) (B). Each point is one individual. Standard length measurements were not obtained for the socialised shallow cohort. The appearance of xanthophore-like cells and iridophores in the fin are unaffected by timing of isolation. In embryo cohorts, xanthophore-like cells first appear in the fin 2-9 days after isolation for imaging (12-19 dpf), and are observed on the initial day of imaging in the yolk closure and socialised cohorts, indicating a likely earlier appearance before imaging began for all individuals. Although the standard lengths at the point of xanthophore appearance were significantly different between all cohorts (Table S2), this simply reflects the different developmental stages at the start of each imaging series, given we infer earlier appearances for yolk and socialised cohorts. Iridophores appear around yolk closure: in embryo cohorts, iridophores first appear in the fin 9-10 days after isolation for imaging (19-20 dpf) and 0-3 days for yolk closure cohorts (20-23 dpf) (Table S2). Most of the socialised cohorts (n=19 out of 22) had iridophores visible in the fin on the initial imaging day, again likely reflecting an earlier undocumented appearance (Table S2). When iridophores are first observed, the socialised cohort standard lengths are significantly different to the embryo and yolk-closure cohorts, again reflecting the different starting developmental stage for this cohort. More informatively, the embryo and yolk-closure cohort standard lengths are not significantly different from each other despite different days since isolation, confirming that the 19-23 dpf (around yolk closure) is the developmental stage at which iridophores appear in the fin across cohorts. Therefore the timing of the first two events are better predicted by days while the later two events are better predicted by standard length (main text), indicating that only iridophore appearance between fin rays 4-5 and formation of iridophore-xanthophore associations are plastic with respect to territoriality timing.

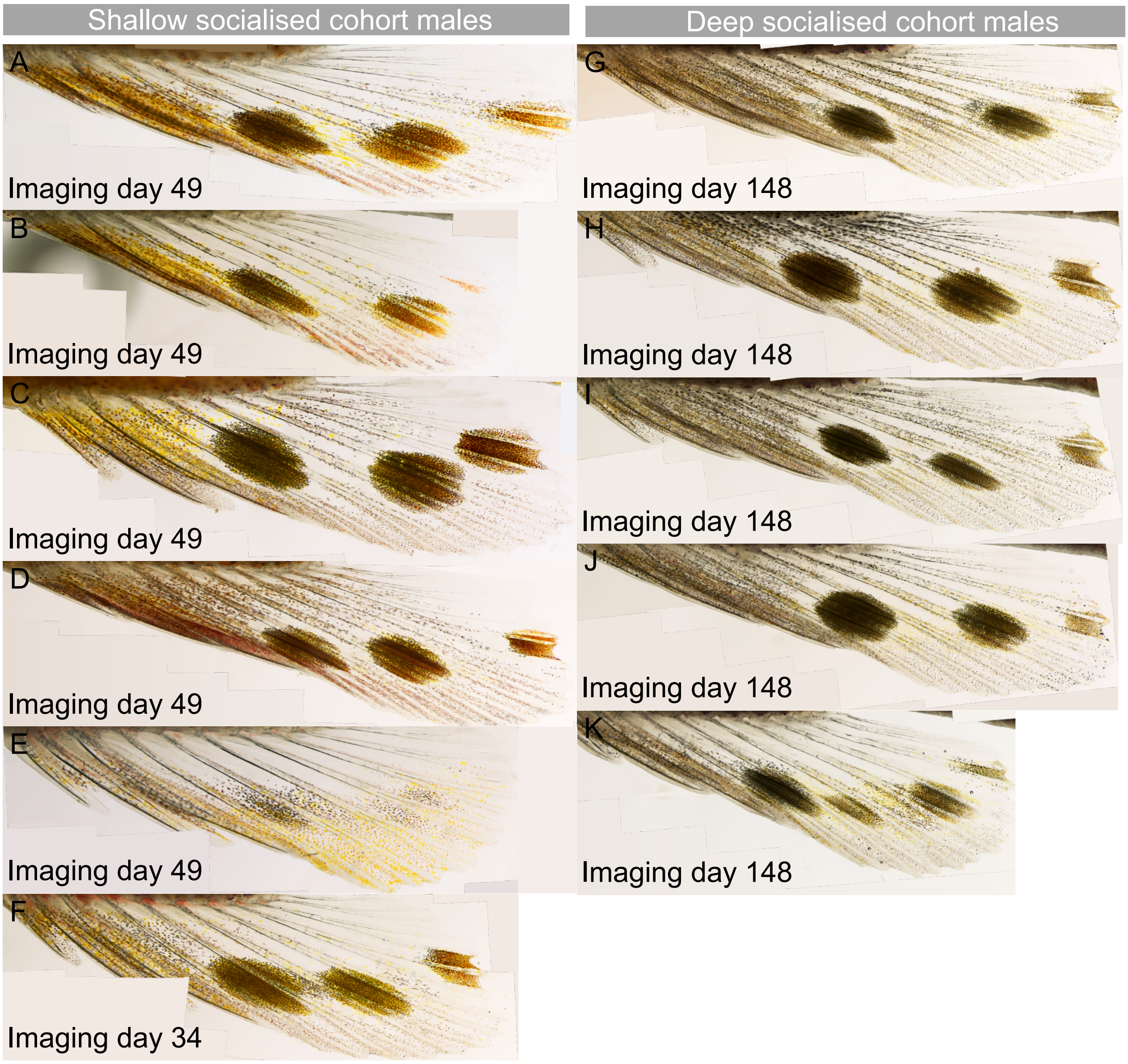


**Fig. S10. Final appearances of male fins in socialised cohorts.** All images were taken on the last day each individual was imaged. (A-F) Shallow socialised cohort. (G-K) Deep socialised cohort. All males have three mature egg-spots except (B) which has two and (E) which has none. The aggregation between the first and second egg-spots in (K) was not considered for positional counting in Table S3 and Figure 6.

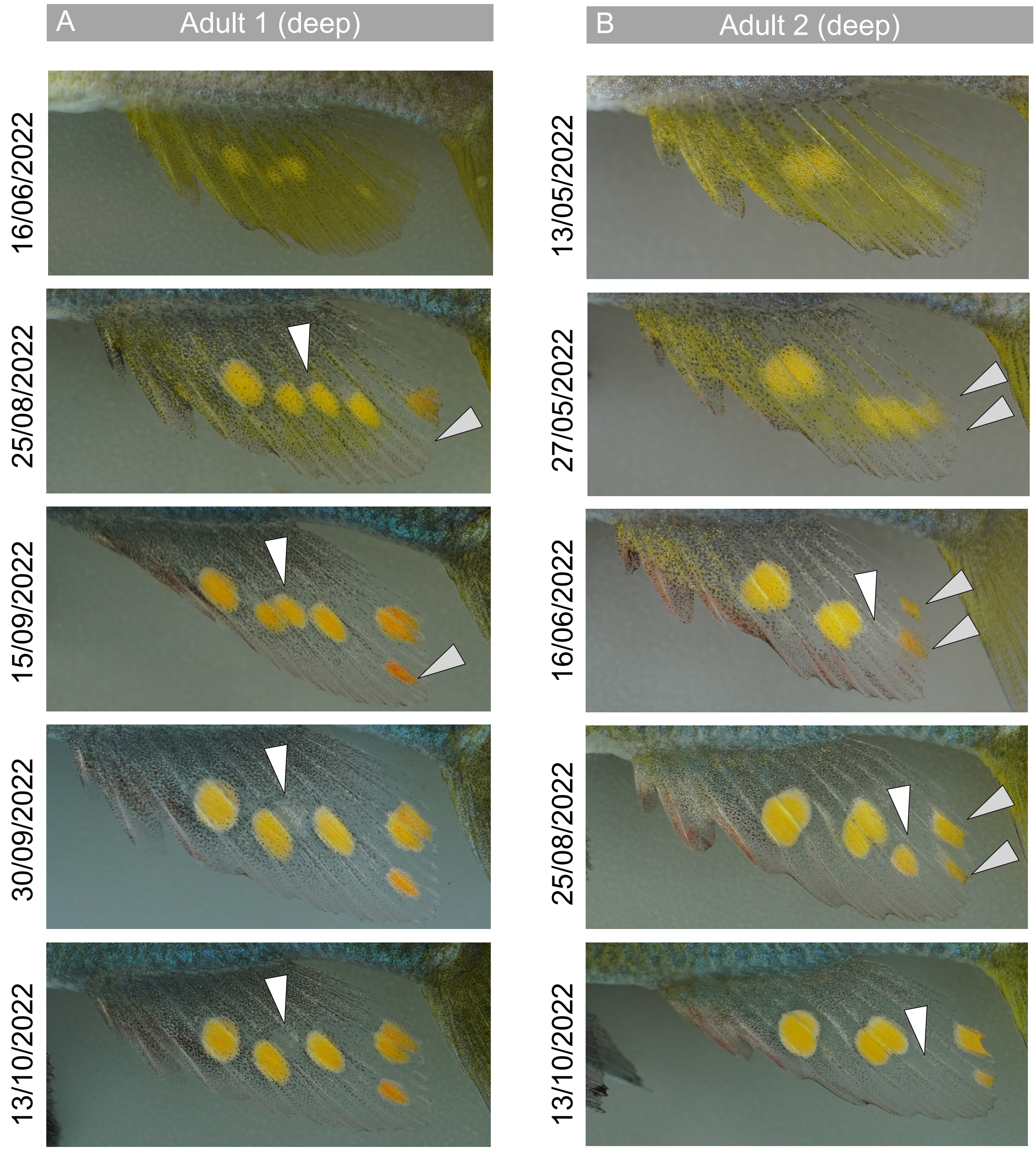


**Fig. S11. Changes in adult egg-spots.** (A,B*)* Photographs of deep morph males isolated with visible aggregations. Dates on which photographs were taken are displayed on the side of each. (A) Adjacent egg-spots can merge (white arrows) and additional egg-spots can form in areas of the fin previously unoccupied by any aggregation or mature egg-spot (grey arrows). (B) Egg-spots can appear and then disappear again in the course of months (white arrows) and large aggregations can develop into two nearby egg-spots which change in size relative to each other (grey arrows).

**Table S1. Primer sequences for the sex determination assays.**

| Primer | PrimerPartner | Sex determination system | Sequence | Tm | GC | Size bp |
|---|---|---|---|---|---|---|
| dup_fwd | dup_rev | *Chromosome 7 gsdf duplication* | TGTCGCGTCATAACGAGGAG | 59.9 | 55 | 207 |
| dup_rev | dup_fwd | *Chromosome 7 gsdf duplication* | AGCTGATCTGGTCCCTCACT | 60 | 55 | |
| control_fwd | control_rev | *Chromosome 7 gsdf duplication* | GCTGCCCACCTCGTAGTAAT | 59.5 | 55 | 402 |
| control_rev | control_fwd | *Chromosome 7 gsdf duplication* | GCACGAGTGGGAACCAGTAA | 60 | 55 | |
| dup_fwd | control_rev | *Chromosome 7 gsdf duplication* | | | | 614 |
| chr19_fwd | chr19_rev | Chromosome 19 TE insertion | AGTCTGGATCGAAGTCCCGT | 60.3 | 55 | 374 |
| chr19_rev | | Chromosome 19 TE insertion | GCAACCACTCTGAAAGTCTCCA | 60.5 | 20 | |
| chr19_sex_ins_fwd | chr19_rev | Chromosome 19 TE insertion | GGGCCTTATTGAATGACAGAGTT | 58.7 | 44 | 248 |
| chr7_fwd | chr7_rev | Chromosome 7 TE insertion | TGGCCTCGGGCATTACTTATT | 60 | 48 | 448 |
| chr7_rev | | Chromosome 7 TE insertion | TTCACTGAACGACATCCTGGT | 59.3 | 48 | |
| chr7_sex3_ins_rev | chr7_fwd | Chromosome 7 TE insertion | AGGTTCTTTGACGTATGATGGT | 57.5 | 41 | 203 |

**Table S2. Results of comparisons between cohorts for timing of early aggregation events, measured by days since initial imaging session or standard length (mm).** Choice of statistical test was determined by number of comparisons: Kruskal-Wallis when only two groups were being compared, Pairwise Wilcox when multiple groups, with Bonferroni adjustment for multiple testing. R code is available on github. Embryos are the only cohort with significantly different days to other cohorts at the point of xanthophores appearing in the fin while all cohorts' standard lengths are significantly different from each other, reflecting that in juvenile cohorts xanthophores are already present in the fin and are recorded on the initial imaging day. Similarly, for iridophore appearance in the fin, embryos are the only cohort with significantly different days, while the standard lengths of embryos and yolk closure cohorts are significantly different from the socialised cohort but not from each other indicating that yolk closure is the stage near which iridophores appear in the fin across cohorts.

When socialised shallow individuals are excluded due to individuals being selected with existing aggregations, for both the appearance of iridophores between the fourth and fifth fin rays and the appearance of associations between iridophores and xanthophores, the days since initial imaging sessions are not significantly different between yolk closure and socialised cohorts while the standard lengths are significantly different.

There is no significant difference between yolk closure and socialised cohorts for the number of days elapsed between iridophore appearance between the 4th and the 5th fin rays and then the  appearance of associations between iridophores and xanthophore-like cells. As such, the duration of egg-spot initialisation was unaffected by the developmental stage at which isolation commenced.

| Event | Measurement | Group 1 | Group 2 | p value | p<0.05? | Test | Division by |
|---|---|---|---|---|---|---|---|

| | | | | | | | |
|---|---|---|---|---|---|---|---|
| Xanthophores in fin | Day | Embryo | Yolk | 7.50E-07 | Yes | Pairwise Wilcox test with bonferroni adjustment | Cohort |
| Xanthophores in fin | Day | Yolk | Socialised | 0.71 | No | Pairwise Wilcox test with bonferroni adjustment | Cohort |
| Xanthophores in fin | Day | Socialised | Embryo | 1.00E-08 | Yes | Pairwise Wilcox test with bonferroni adjustment | Cohort |
| Xanthophores in fin | SL | Embryo | Yolk | 0.00029 | Yes | Pairwise Wilcox test with bonferroni adjustment | Cohort |
| Xanthophores in fin | SL | Yolk | Socialised | 0.00019 | Yes | Pairwise Wilcox test with bonferroni adjustment | Cohort |
| Xanthophores in fin | SL | Socialised | Embryo | 0.00019 | Yes | Pairwise Wilcox test with bonferroni adjustment | Cohort |
| Iridophores in fin | Day | Embryo | Yolk | 2.50E-05 | Yes | Pairwise Wilcox test with bonferroni adjustment | Cohort |
| Iridophores in fin | Day | Yolk | Socialised | 0.38 | No | Pairwise Wilcox test with bonferroni adjustment | Cohort |
| Iridophores in fin | Day | Socialised | Embryo | 3.40E-06 | Yes | Pairwise Wilcox test with bonferroni adjustment | Cohort |
| Iridophores in fin | SL | Embryo | Yolk | 1 | No | Pairwise Wilcox test with bonferroni adjustment | Cohort |
| Iridophores in fin | SL | Yolk | Socialised | 0.00019 | Yes | Pairwise Wilcox test with bonferroni adjustment | Cohort |
| Iridophores in fin | SL | Socialised | Embryo | 0.00037 | Yes | Pairwise Wilcox test with bonferroni adjustment | Cohort |
| Iridophores 4-5 | Day | Yolk deep | Socialised deep | 1 | No | Pairwise Wilcox test with bonferroni adjustment | Cohort and morph |
| Iridophores 4-5 | Day | Yolk shallow | Socialised deep | 1 | No | Pairwise Wilcox test with bonferroni adjustment | Cohort and morph |
| Iridophores 4-5 | Standard length | Yolk | Socialised (deep only SL) | 5.55E-05 | Yes | Kruskal-Wallis rank sum test | Cohort |

| | | | | | | | |
|---|---|---|---|---|---|---|---|
| Associations | Day | Yolk deep | Socialised deep | 1 | No | Pairwise Wilcox test with bonferroni adjustment | Cohort and morph |
| Associations | Day | Yolk shallow | Socialised deep | 0.389 | No | Pairwise Wilcox test with bonferroni adjustment | Cohort and morph |
| Associations | Standard length | Yolk | Socialised (deep only SL) | 4.43E-04 | Yes | Kruskal-Wallis rank sum test | Cohort |
| Time between Iridophores 4-5 and Associations | Days | Yolk shallow | Socialised deep | 0.5242 | No | Pairwise Wilcox test with bonferroni adjustment | Cohort and morph |
| Time between Iridophores 4-5 and Associations | Days | Yolk deep | Socialised deep | 0.9636 | No | Pairwise Wilcox test with bonferroni adjustment | Cohort and morph |
| Time between Iridophores 4-5 and Associations | Days | Yolk shallow | Yolk deep | 0.9909 | No | Pairwise Wilcox test with bonferroni adjustment | Cohort and morph |

**Table S3. Variability in subsequent egg-spot formation.** Proportion of male individuals from socialised cohorts having an egg-spot at each position. Positions are described by the fin rays that the aggregations are bounded by at either side. Positions are measured on the final imaging session for mature male juveniles (see figure S10). Note that one male did not develop a third spot by the final imaging session. Only mature egg-spots with dense and defined cellular aggregations were considered. Data is represented in a diagram in Figure 6a.

| Position (fin rays bounded by) | Number | Proportion (%) |
|---|---|---|
| First spot (10/10 individuals) | | |
| 4-6 | 3 | 30 |
| 3-6 | 6 | 60 |
| 4-5 | 1 | 10 |
| Second spot (10/10 individuals) | | |
| 5b to 7b | 2 | 20 |
| 6a to 8a | 1 | 10 |
| 5b to 8a | 1 | 10 |
| 6 to 7b | 1 | 10 |
| 6b to 8b | 3 | 30 |
| 6a to 8b | 1 | 10 |
| 7a to 8b | 1 | 10 |

| Third spot (9/10 individuals) | | |
|---|---|---|
| 7b to 9a | 1 | 11.11 |
| 8b to 9b | 2 | 22.22 |
| 8a to 9b | 1 | 11.11 |
| 8a to 9a | 1 | 11.11 |
| 9a to 10 | 2 | 22.22 |
| 8b to 10 | 2 | 11.11 |
